# Adaptive and maladaptive genetic diversity in small populations; insights from the Brook Charr (*Salvelinus fontinalis)* case study

**DOI:** 10.1101/660621

**Authors:** Anne-Laure Ferchaud, Maeva Leitwein, Martin Laporte, Damien Boivin-Delisle, Bérénice Bougas, Cécilia Hernandez, Éric Normandeau, Isabel Thibault, Louis Bernatchez

## Abstract

Investigating the relative importance of neutral *versus* selective processes governing the accumulation of genetic variants is a key goal in evolutionary biology. This is particularly true in the context of small populations, where genetic drift can counteract the effect of selection. In this study, we investigated the accumulation of putatively beneficial and harmful variations using 7,950 high-quality filtered SNPs among 36 lacustrine, seven riverine and seven anadromous Brook Charr (*Salvelinus fontinalis*) populations (n = 1,193) from Québec, Canada. Using the Provean algorithm, we observed an accumulation of deleterious mutations that tend to be more prevalent in isolated lacustrine and riverine populations than the more connected anadromous populations. In addition, the absence of correlation between the occurrence of putative beneficial nor deleterious mutations and local recombination rate supports the hypothesis that genetic drift might be the main driver of the accumulation of such variants. Despite the effect of pronounced genetic drift and limited gene flow in non-anadromous populations, several loci representing biological functions of potential adaptive significance were associated with environmental variables, and particularly with temperature. We also identified genomic regions associated with anadromy. We also observed an overrepresentation of transposable elements associated with variation in environmental variables, thus supporting the importance of transposable elements in adaptation.

## Introduction

The fate of wild populations exposed to environmental variation is determined by an interplay between genetic variation and demography (Lande 1988). Genome carries beneficial but also deleterious mutations (between 500 and 1,200 deleterious variants have been estimated in a single human genome (Fay et al 2001; Sunyaev et al 2001)) potentially affecting individual fitness. A lack of genetic diversity has been implicated in many populations and species extinction since it increases the chances of recessive deleterious alleles to be expressed and decreases the chances for an individual to have genotype matching the environmental challenges (Fagan and Holmes 2006; Lynch, Conery and Buerger, 1995). The potential negative impact that inbreeding will have on population health and reproduction compared to an outbred population is referred to as “genetic load” (Crow, 2001). According to population genetics theory, the accumulation of deleterious mutations and therefore the genetic load depends on multiple factors such as effective population size, admixture and selection (Whitlock and Buerger 2004). Although natural selection may increase the frequency of adaptive mutations and remove deleterious alleles at the same time (Whitlock et al 2003), deleterious variants may also hitchhike with advantageous mutations to high frequencies if they have intermediate fitness costs (Chun and Fay 2011). Additionally, recombination facilitates the removal of maladaptive alleles from a population (Felsenstein 1974), and genomic regions of low recombination are more likely to accumulate deleterious alleles as the efficiency of selection is reduced (Charlesworth 2009). However, systems in which stochastic demographic processes, (*e.g.* genetic drift, bottlenecks) prevail over natural selection (*e.g.* small isolated populations) may allow deleterious variants to increase in frequency (Lynch, Conery and Buerger 1995) and inbreeding can unmask such recessive deleterious alleles affecting fitness of individuals and populations. Thus, at a genomic level, when demographic factors are mainly responsible for the accumulation of deleterious mutations, a random genome-wide distribution of deleterious variants is expected. In turn, when selection is mainly responsible for the accumulation of those variants, maladaptive variants are expected to accumulate mostly in low recombination rate regions.

Anthropogenic climate change induced an elevation of global temperatures (Kerr 2007; Marcott et al. 2013; Moss et al. 2010), and its effects on plants and animals are already evident, especially for ectothermic organisms, and particularly so in northern latitudes (Paaijmans et al 2013). Almost all fishes are ectothermic and thereby unable to adjust body temperature physiologically, which implies that their metabolism, reproduction and growth are strongly influenced by temperature (Dickson and Graham 2004). Thermal tolerance is known to be partly genetically based in fish (*e.g.* Dammark et al 2018). Research on temperature adaptation and climate change adaptability in fishes has focused on local or environment-dependent temperature related adaptation in life-history traits (Bradbury et al 2010) particularly in salmonids fish (Narum et al 2010; Hecht et al 2015).

The Brook Charr (*Salvelinus fontinalis*, Mitchill) is an endemic salmonid of eastern North America (Scott and Crossman 1973) and ranks among the most highly structured animal species (Gyllensten 1985; Ward, Woodwark, and Skibinski 1994), with most of its genetic variance partitioned among major drainages (Ferguson et al 1991; Perkins et al 1993; Danzmann and Ihssen 1995; Angers and Bernatchez 1998). Previous phylogeographic studies revealed that a single glacial race of Brook Charr colonized most of Northeastern America (Ferguson, Danzmann, and Hutchings 1991; Jones, Clay, and Danzmann 1996). Very low effective population sizes have been reported for this species, particularly so in lacustrine populations that are generally isolated with limited gene flow between them (Gossieaux et al 2019; median Ne for the studied populations here is 35, IC 32:38, data not shown). Besides resident populations, many Brook Charr populations are anadromous. Anadromy, a trait shared by many salmonid species, refers to a migratory life cycle whereby individuals are born in fresh water, feed and grow in salt water and return to fresh water to reproduce. Consequently, these populations are more likely to be connected by gene flow than resident ones (Castric and Bernatchez 2003). Previous studies showed that a significant proportion of phenotypic variation in migratory traits has a genetic basis (Liedvogel et al 2011, including in Brook Charr (Boulet et al. 2012). A common genetic mechanism was revealed to be involved in the life-history transition from anadromy to residency in the Rainbow and Steelhead Trout (*Oncorhynchus mykiss*, Hecht et al 2012; Pearse et al 2014). Freshwater migration length and elevation steepness are known to have a strong effect on the bioenergetics costs of migration, including salmonids (Bernatchez & Dodson, 1987), which can drive local adaptation of life history traits (Pearse et al 2014, Moore et al 2017), including in Brook Charr (Fraser and Bernatchez (2005); Crespel et al 2017).

In this study, we investigate the importance of demographic processes *versus* selection governing the accumulation of both adaptive and maladaptive (deleterious) mutations in the Brook Charr (*Salvelinus fontinalis*, Mitchill). By adaptive, we refer to variations linked to environmental variables, hypothesizing that some mutations are more beneficial in one environment than in another, despite no fitness values corroborate it; and by maladaptive, we refer to variations having putative damaging effect on the individual fitness inferred from the known functionality of a given mutation (see below). More specifically, based on a GBS dataset from samples of lakes, rivers and anadromous Brook Charr locations distributed all throughout the province of Quebec (Canada), we aim to (i) assess the extent of genetic diversity and differentiation among localities and habitats, (ii) detect the presence of putative adaptive mutations associated to environmental variables as well as maladaptive mutations and (iii) estimate the recombination rate along the genome and its potential role in explaining the genomewide distribution of those mutations. Finally, we combine all results in order to explore the relative roles of neutral and selective processes on the accumulation of maladaptive mutations among salmonids populations. We predict that if limited gene flow and the occurrence of pronounced genetic drift are the main factors governing the accumulation of deleterious variants, we expect to observe a lower frequency of deleterious mutations among potentially more connected anadromous and river populations compared to unconnected lacustrine populations. If selection is the main factor explaining mutation load we expect to observe a higher accumulation of deleterious mutations in genomic regions of low recombination rate, as well as potentially observing a positive relationship between the accumulation of deleterious mutations and inferred recombination rate.

## Methods

### Sampling and study system

A total of 1,193 individuals sampled from 50 sites (36 lakes, 7 rivers and 7 anadromous populations) from 2014 to 2015 (for lakes and rivers) and from 2000 to 2001 for anadromous sites (Castric and Bernatchez 2003) in Québec, Canada, were successfully genotyped (Figure 1, Table 1). The median number of individuals per site was 24 (Table 1). Fish were sampled by technicians of the Ministère des Forêts, de la Faune et des Parcs du Québec (MFFP) and the Société des Établissements de Plein Air du Québec (SEPAQ) using gillnets or by anglers using fishing rods. Adipose fin clips were preserved in 95% ethanol. Populations were chosen posterior to sampling according to their geographic location, the documented absence of stocking, and the availability of tissues from which good quality DNA could be extracted (A_260_/A_280_ ratio between 1.7 and 2.0, and high molecular weight together with no smears while migrated on an agarose gel). The size of the lakes sampled ranged from 3 to 5591 hectares, with a median value of 66 hectares (Table 1). Latitude ranged from 45.378 to 57.918 and longitude from −79.194 to −57.949. Records of water temperature were unavailable for most of the sites and so records of air temperature were used to estimate 10-year averages (2004-2015) of minimum, maximum, mean minimum, mean maximum and mean air temperature for each lake from the database available in BioSim (Réginiére *et al*. 2014). Air temperature has been shown to be linearly correlated with growth rate in *Salvelinus* and thus is a good variable to study the role of temperature in shaping local adaptation (Black et al. 2013; Torvinen 2017; Perrier, Ferchaud et al. 2017; Ferchaud, Laporte et al. 2018). Annual average air temperature ranged from −5.04 to 6.29 °C (Figure 1, Table 1). Other environmental variables of interest were gathered using BioSim to get 10-year averages of dew point, frost free days, total of radiation and Growing Degree-Day (GDD) was provided by Ministère des Forêts, de la Faune et des Parcs (MFFP) du Québec. Briefly, the basic concept of GDD is that development will only occur if the temperature exceeds some minimum development threshold, or base temperature (TBASE). The base temperatures are determined experimentally and are different for each organism. To calculate GDDs, one must (i) estimate the mean temperature for the day (by adding together the high and low temperature for the day and dividing by two); (ii) compare the mean temperature to the TBASE. Then, if the mean temperature is at or below TBASE, the Growing Degree Day value will be zero. If the mean temperature is above TBASE, the Growing Degree Day amount will be equal to the mean temperature minus TBASE. For example, if the mean temperature was 25° C, then the GDD amount equals 10 for a TBASE of 15° C. For each locality, we averaged GDD values collected over a 10-year period before the sampling (2004-2015) using a base temperature of 5°C (temperature usually employed in such geographic regions, MFFP personal communication). All the environmental variables are reported in Supplementary Table 1.

**Figure 1.**
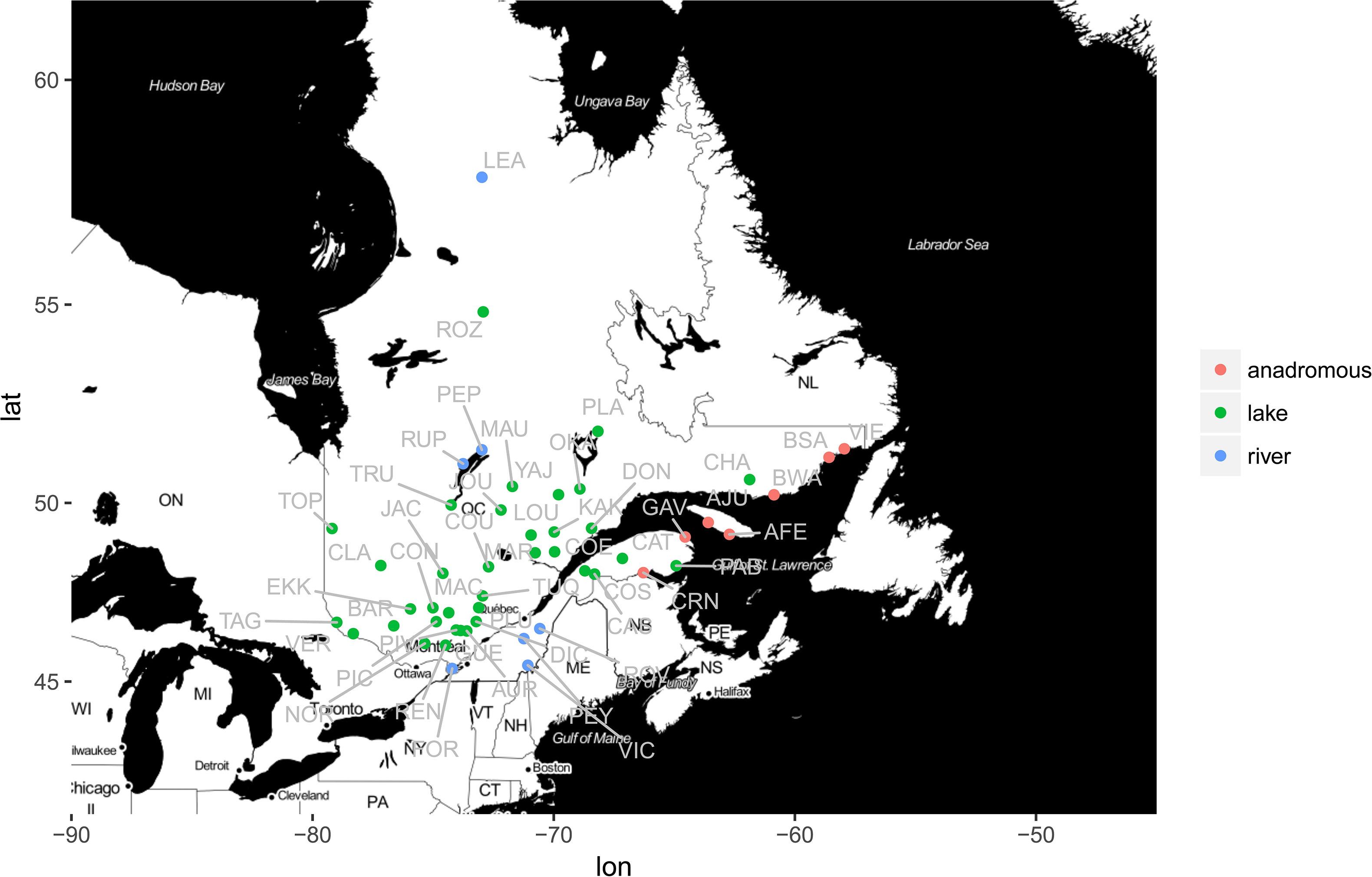
Geographic location of the 50 sampling sites studied across the province of Quebec, Canada; 36 lakes in green, 7 rivers (in blue) and 7 anadromous localities (in red). Each locality is labelled using a three-letter code displayed in grey. The map drawn using the R-package ggmap (Kahle & Wickham 2013) with R 3.1.0 (R development core team 2017).

**Table 1:**
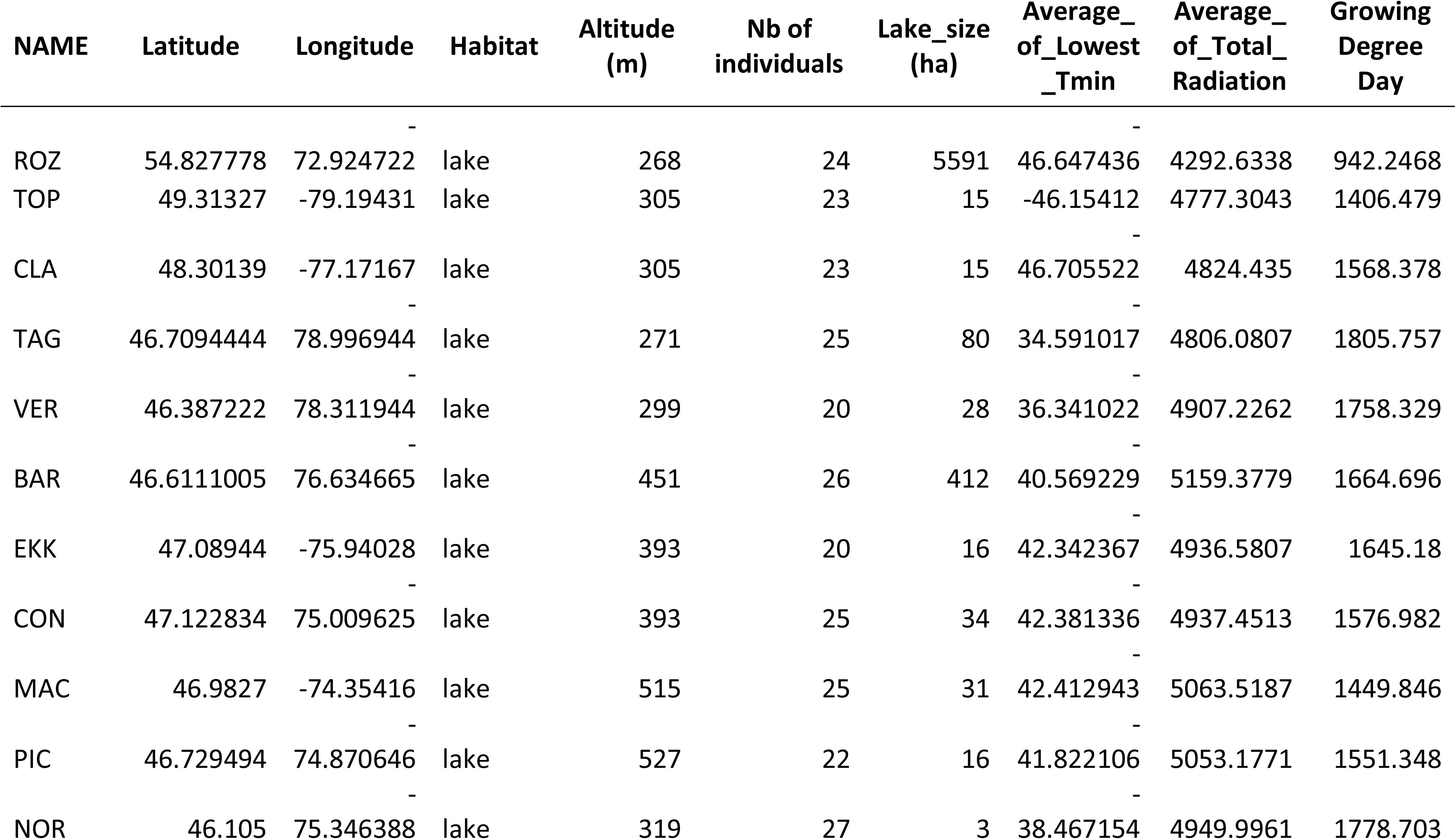
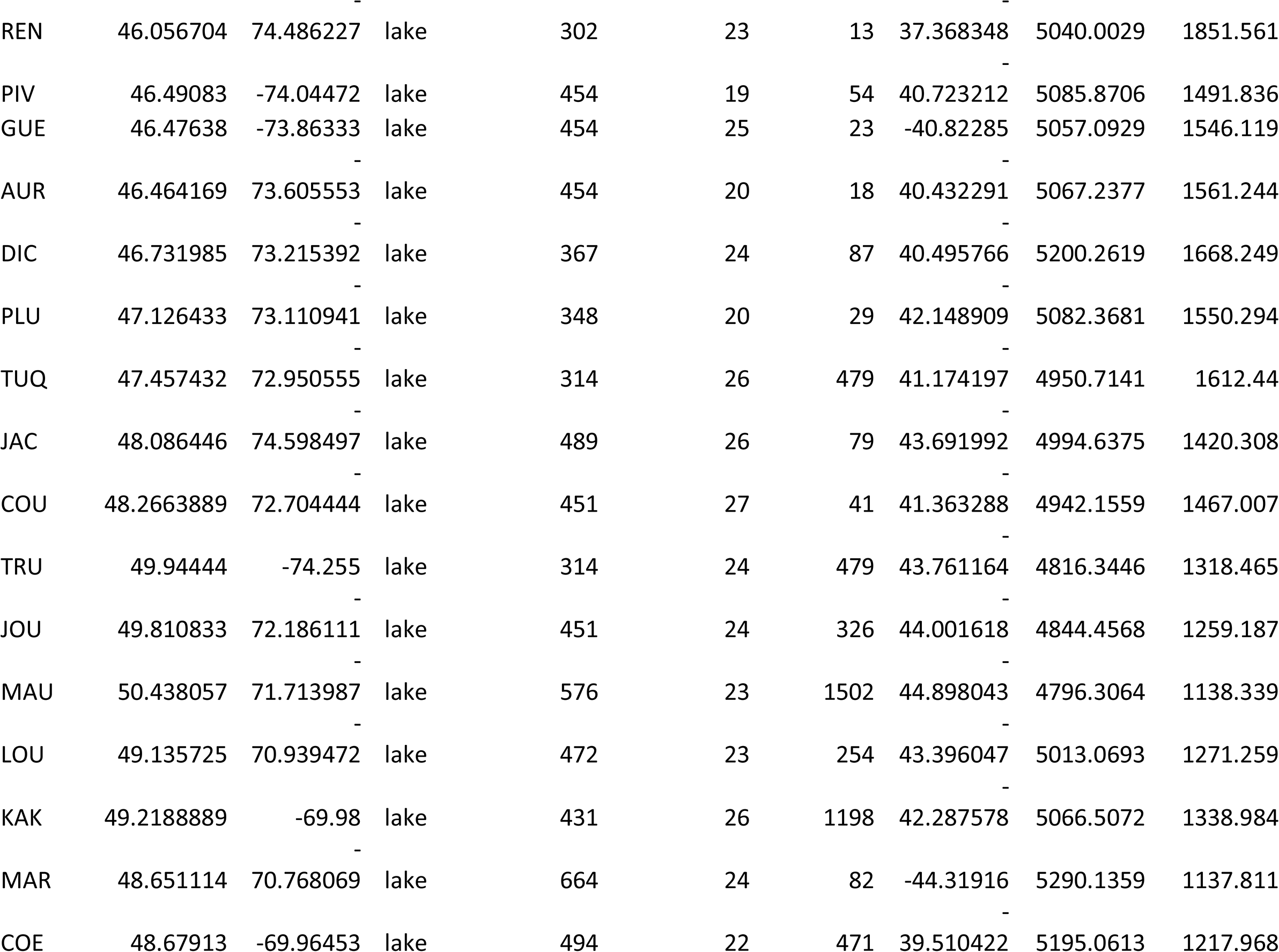
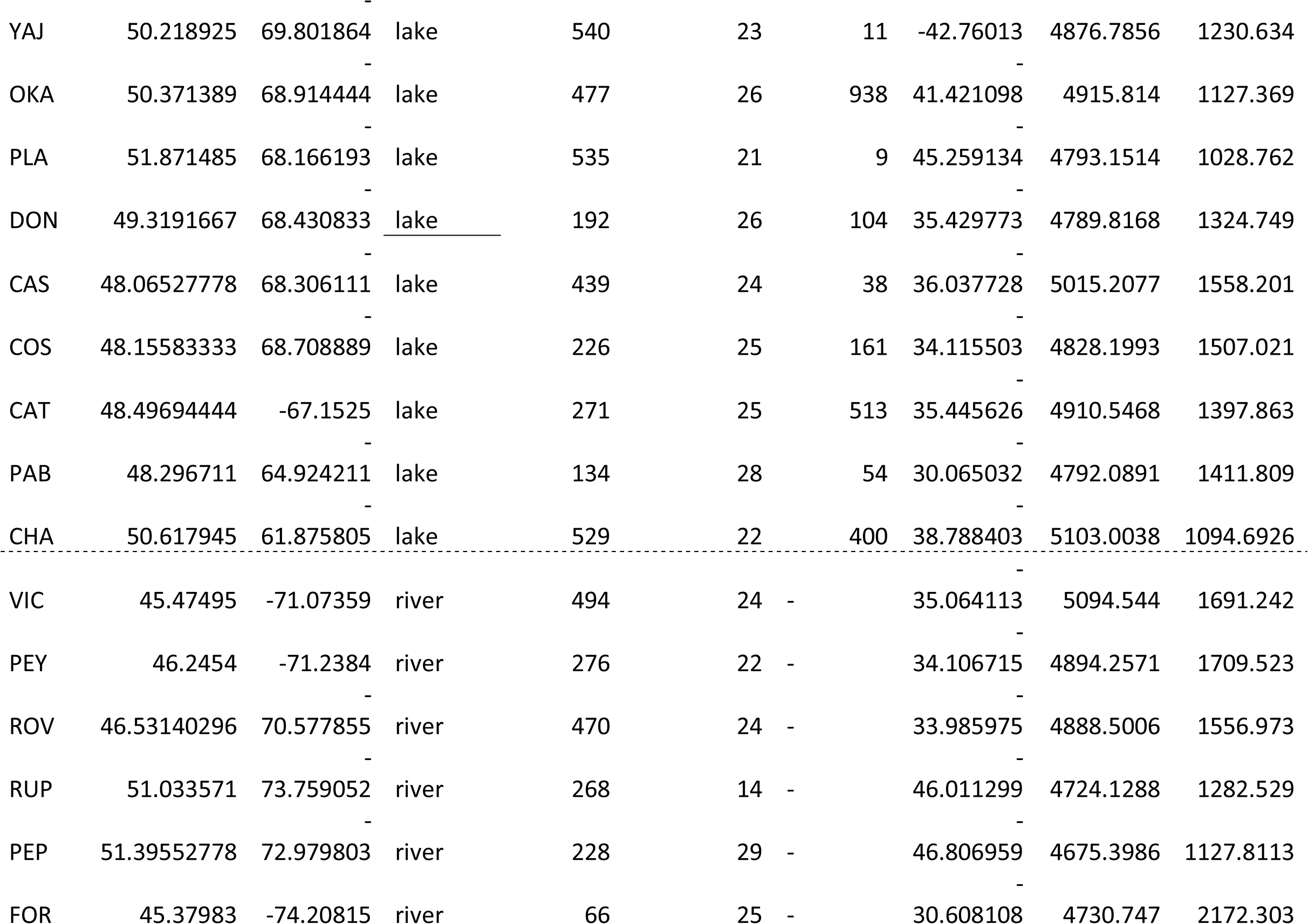
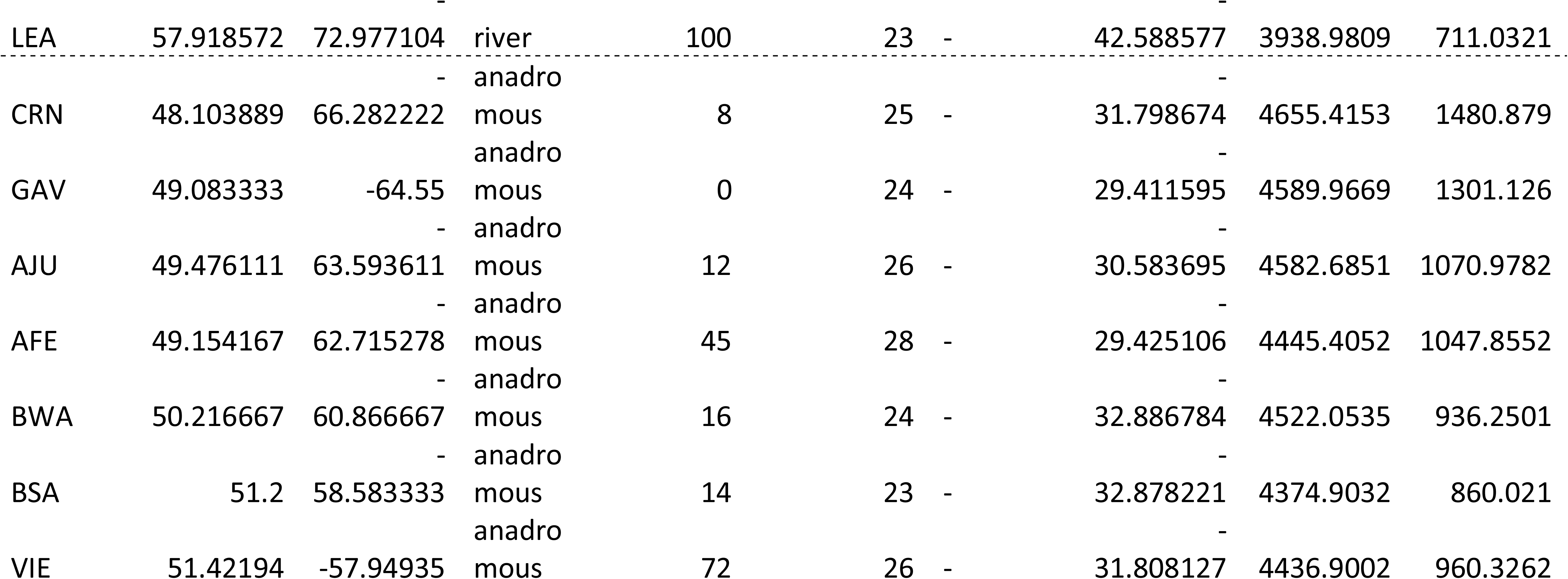
Sampling locations and corresponding sample size, latitude, longitude, habitat type, altitude, lake size average of minimum air temperatures, average of total radiation and Growing Degree-Day. The entire dataset of environmental variables used in this study are reported in Supplementary Table 1. Localities are ordered to their habitat type and subsequently according to geography (mainly from West to East), same order is used all along the figures and tables of the manuscript.

### Molecular analyses

Total DNA was extracted from adipose fin tissue (5 mm^2^) using a slightly modified version of Aljanabi and Martinez (1997) salt extraction protocol. Sample concentration and quality were checked using 1% agarose gel and a NanoDrop 2000 spectrophotometer (Thermo Scientific). DNA was quantified using the PicoGreen assay (Fluoroskan, Ascent FL; Thermo Labsystems). Genomic DNA was normalized to obtain 20 ng/μl in 10 μl (200 ng) for each individual. The sequencing libraries were created accordingly to Mascher et al (2013) protocol. Namely, in each sample, a digestion buffer (NEB4) and two restriction enzymes (*PstI* and *MspI*) were added. After a two-hour incubation period at 37°C, enzymes were inactivated by a 20-min incubation period at 65°C. Then, the ligation of two adaptors was performed using a ligation master mix followed by the addition of T4 ligase and completed for each sample at 22°C for 2 hr. Enzymes were again inactivated by a 20-min incubation period at 65°C. Finally, samples were pooled in 96-plex and QIAquick PCR purification kit was used to clean and purified the DNA. After library PCR amplification, sequencing was performed on the Ion Torrent Proton P1v2 chip at the genomic analysis platform of IBIS (Institut de Biologie Intégrative et des Systèmes, Université Laval).

### Bioinformatics

The program FastQC 0.11.1 (Andrews 2010) was used to check raw reads for overall quality and presence of adapters. All the bioinformatic steps, options, and software versions employed in the subsequent GBS pipeline are detailed in Supplementary Table 2. Briefly, we used cutadapt v1.8.1 (Martin 2011) to remove the adapter from raw sequences and STACKS v1.40 *process_radtags* to demultiplex the samples and do the quality trimming (Catchen et al 2013). Stacks were created using *ustacks* allowing a maximum of two nucleotide mismatches (M = 2) among primary reads and four nucleotide mismatches (N = 4) among secondary reads. No polymorphisms were called if they were only present in the secondary reads. M and N values were identified as an optimum threshold according to preliminary tests across different values of those parameters according to Paris *et al*. (2017). *Cstacks* was used to build a reference catalog with all loci identified across a subset of 204 individuals corresponding to the four individuals having the highest number of reads in each sampling site. The minimum stacks depth was set to four (m = 4). Sets of stacks were searched against the catalog using *sstacks*. *rx-stacks* was used to correct genotypes and *c-stacks* and *s-stacks* were run again with the correction. The stacks’ module POPULATIONS was then used to call genotypes. Only individuals with less than 40% of missing genotypes were considered for the subsequent analysis and the R package *stackr* (Gosselin 2017a) was used to remove loci with more than two alleles. Output file was also filtered using custom script to retain high-quality SNPs (available at https://github.com/enormandeau/stacks_workflow). Low variant loci were removed (minor allele frequency <0.05), and a dataset was built keeping only a single SNP per locus (the one harboring the highest minor allele frequency) to avoid linkage disequilibrium bias (see Supplementary Table 2 for details about every step). Finally, the function detect_het_outliers of the R package *radiator* (Gosselin 2017b) was run to estimate the overall genotyping error rate and the overall miscall rate of the final dataset.

### Population differentiation

A Principal Component Analysis (PCA) was performed using the function glPCA() of adegenet 2.1.2 package in R to seek a summary of the diversity among all localities. The R function ggplot2 (Wickham 2009) was then used to plot the results of this PCA according (i) the sampling locality origin of each sample and (ii) the habitat type (anadromous, lake or river). Genetic structure among populations was also investigated with ADMIXTURE 1.3.0 program (Alexander, Novembre and Lange 2009) for K ranging from 2 to 60 across all localities and subsequently from 2 to 14 across anadromous localities only. The Cross-Validation indices were used to discuss the best values of K across all localities and across anadromous localities. Both for PCA and Admixture analysis, the dataset with only a single SNP per locus was employed and converted respectively in genlight object or plink file. The extent of genetic differentiation among sampling locations was computed by average F_ST_ estimations with Genodive 2.0.27 (Meirmans and Tienderen 2004) among (i) all locality pairs and then considering the three habitat types separately, (ii) among pairs of anadromous locations (iii) among pairs of lacustrine locations and (iv) among pairs of riverine locations. Both the pairwise population differentiation (*F*_ST_) and the mean pairwise *F*_ST_ per population were reported. Finally, signals of isolation-by-distance were tested using a correlation between genetic differentiation (Fst/(1-Fst)) and log(geographic distance) (Rousset, 1997) considering (i) all localities altogether and (ii) only lacustrine locations. The correlation between river distance (more representative of riparian fish rather than geographic distance) and genetic differentiation was also explored across three different river drainages (*i.e.* river drainages with more than 2 localities, respectively 4, 9 and 4 localities). Results are not different from those obtained with geographic distance using the full dataset and thus are not shown.

### Population genetic diversity

We documented the extent of genomic diversity within and among populations by estimating the nucleotide diversity “Π”. Π was estimated within each individual and then averaged by population, on all sequenced nucleotides including variant and non-variant sites (= 34 789 nucleotides), using the function summary haplotypes from stackr, R-package (Gosselin 2017a). The ratio of polymorphic SNPs was reported for each population as well as the observed heterozygosity estimated with Genodive. t-tests were performed in order to test if the genetic diversity (Π) was significantly different among the three habitat types. Furthermore, using linear models in R, correlations between average Π or average pairwise *F*_ST_ were determined either against latitude and altitude.

### Proportions of maladaptive (deleterious) mutations

To identify putative deleterious mutations, all defined loci (80 bp long) were first used in a BLAST query against the Rainbow Trout (*Oncorhynchus mykiss*) transcriptome (Berthelot et al., 2014) using blastx. All hits that had higher than 25 amino acid similarity and more than 90% similarity between the query read and the transcriptome sequence were retained. Both variants of each SNP in each query read were used with BLAST against the transcriptome, and translation results were compared pairwise. For each locus, results were only kept if the lowest e-value hit for both variants was the same (i.e., same protein name and length), and these were used to extract the protein sequence and identity for the following step. Finally, PROVEAN (protein variation effect analyser; Choi et al 2012) was used to predict the deleterious effect of nonsynonymous mutations with the default deleterious threshold value (−2.5), commonly employed in other studies (Renault and Rieseberg 2015, Perrier, Ferchaud et al 2017, Ferchaud et al 2018). This program uses a versatile alignment-based score to predict the damaging effects of variations not limited to single amino substitutions but also in-frame insertions, deletions and multiple amino acid substitutions. This alignment-based score measures the change in sequence similarity of a query sequence to a protein sequence homolog before and after the introduction of an amino acid variation to the query sequence. Results of PROVEAN have previously been compared in another salmonid (Lake Trout) to another program PolyPhen-2 and results inferred from the two methods were highly congruent (Ferchaud, Laporte et al 2018). For variant harboring a putative deleterious mutation, we hypothesize that the minor allele is the one having a deleterious effect (since it is present in lower frequency at a global level). Because deleterious alleles are selected against, theory predicts that their frequencies should be lower than non-deleterious ones (Fay et al 2001). Thus, *t* tests were also used to verify whether average “deleterious allele frequency” was lower than the ones of “nonsynonymous, but non-deleterious SNPs” and to that of “anonymous” (SNPs that did not mapped against the genome) and “synonymous SNPs” within each habitat type. Next, the proportion of deleterious mutations (defined as the number of SNPs showing a deleterious mutation in a given population over the number of SNPs harboring a deleterious mutation across all populations) and the ratio of the proportion of deleterious mutations over the proportion of polymorphic SNPs were estimated and compared among populations. The latter ratio is expected to be lower in populations that purge deleterious mutations faster (see Perrier, Ferchaud et al 2017) and was compared between habitat types using a *t* test.

### Putative adaptive mutations: gene-environment associations

Redundancy analysis (RDA) was conducted as a multi-locus genotype-environment association (GEA) method to detect loci under selection. RDA is an analog of multivariate linear regression analyses, utilizing matrices of dependent and independent (explanatory) variables. The dependent matrix is represented by the genotypic data (here a matrix of minor allelic frequencies by population), and the explanatory variables are comprised into an environmental matrix. Two analyses were performed. First, we aimed to identify SNPs associated to anadromy and then to identify SNPs associated to environmental variables among lacustrine populations.

Forester et al (2018) recommend to not control by spatial autocorrelation for RDA multi-locus outlier identification, however, because the sampled anadromous localities are mainly distributed in the eastern part of our sampling range, we conducted two RDAs, one controlling for spatial autocorrelation and one without controlling for spatial autocorrelation. Then we conservatively retained SNPs that were discovered to be significantly linked to anadromy by both approaches. In order to correct for spatial autocorrelation, a distance-based Moran’s eigen-vector map (db-MEM) based on latitude and longitude, all pre-transformed in meters with the function “*geoXY”* of the R package “SoDA,” was produced. The habitat (anadromous *versus* resident (lakes + rivers)) constituted the explanatory matrix and the db-MEMs related to spatial components constituted the conditioning matrix controlling for autocorrelation. The function “*rda”* was used to compute the RDAs on the model (see Laporte et al 2016; Le Luyer et al 2017; Marengo et al 2017 for examples of similar methodology). An analysis of variance (|ANOVA; 1,000 permutations) was then performed to assess the global significance of the RDAs and the percentage of variance explained (PVE) was computed with the function “*RsquareAdj”.* When not mentioned, R functions were part of the “vegan” package (Oksanen et al 2019). SNPs linked to anadromy were then defined following instructions from the online tutorial proposed by Brenna Forester (Forester et al 2018; https://popgen.nescent.org/2018-03-27_RDA_GEA.html). Only SNPs in common between both RDAs were considered as candidates underlying adaptation to anadromy.

Secondly, we identified SNPs associated with environmental variables across all lakes following the same instructions (https://popgen.nescent.org/2018-03-27_RDA_GEA.html). First, among all environmental variables, we removed variables with correlated predictors (r Pearson > 0.7) and thus kept only Low Minimum Temperature, Total of radiation and the GDD (hereafter respectively named LowTmin, AvRad and GDD) to identify candidate SNPs potentially involved in local adaptation. To identify the best model, the function “*ordistep”* was used to select the best explanatory variables. As recommended by Forester et al (2018), we did not control for spatial autocorrelation among lacustrine locations since simulations showed that this diminishes the power of detection without decreasing false positives in similar demographic scenarios. For subsequent analyses, we removed from all SNPs defined in association with environmental variables those that were simultaneously discovered as putatively deleterious, in order to properly identify putative beneficial mutations.

### Genomic position and gene ontology

Our 4 729 reads were mapped against the Arctic Charr reference genome (Christensen et al 2018) using BWA (Li and Durbin 2009). GWAN (https://github.com/enormandeau/gawn) was performed to annotate the Arctic Charr genome with the Brook Charr transcriptome (Pasquier et al 2016). Since Arctic Charr and Brook Charr differ in their number of chromosomes (32 *versus* 42 linkage groups, respectively), we took advantage of the Brook Charr high density linkage map (Sutherland et al 2016) to retrieve the relative position of our loci (see Leitwein et al. 2017 for the methods). Briefly, we anchored the 3,826 mapped RAD loci of the Brook Charr linkage map (Sutherland et al 2016) to the Arctic Charr reference genome (Christensen et al 2018). This was possible after controlling for synteny and collinearity between the Arctic Charr (Nugent et al 2017) and the Brook Charr linkage map (Sutherland et al 2016) using the MAPCOMP pipeline (Sutherland et al 2016). Then, we were able to create an ordered loci list for each of the 42 Brook Charr linkage groups (*i.e.* we reconstructed a collinear reference genome for the Brook Charr). We particularly recorded the annotations for the putative deleterious mutations and SNPs for which allele frequencies were correlated with anadromy and with temperature and or GGD for lakes (see Results). For doing so, we added 10 000 bp before and after the SNP position to account for physical linkage.

### Effect of recombination rate on the genomewide distribution of putative adaptive and maladaptive mutations

The local Brook Charr recombination rate was estimated by comparing the genetic positions (cM) retrieved from the Brook Charr high density linkage map (Sutherland et al., 2016), to the physical positions (bp) retrieved from our reconstructed Brook Charr reference genome with MAREYMAP (Rezvoy et al 2007). The degree of smoothing was set to 0.9 (*span*) for the polynomial regression method (Loess methods). To infer the recombination rates of the markers not included in the linkage map, we computed the weighted mean recombination rate using the two closest markers based on their relative physical positions. Putative adaptive and maladaptive mutations were plotted along the inferred recombination rate across the 42 Brook Charr linkage groups. Differences in mean recombination rate between types of mutations were tested between markers linked *versus* non-linked markers to environmental variables (*e.g.* linked *versus* non-linked to temperature) using a Wilcoxon test. For the non-synonymous mutations, the relationship between Provean scores and inferred recombination rate was explored using a linear model in R.

## Results

### DNA sequencing and genotyping

The total number of de-multiplexed and cleaned reads was 2,989,523,556 with an average of 2,002,360 reads per individual. After DNA control quality and filtering out individuals with more than 40 % of missing genotypes (the concerned missing data represents 7.1% of the global genotype data set), 1,193 of 1,493 individuals (80%) (with an average number of 2,166,685 reads per individual) were kept for the subsequent analyses. After filtering for quality 7,950 SNPs were kept (distributed in 4,729 loci, Supplementary Table 3). Finally, for the SNPs retained, the overall genotyping error was estimated to be 1e-04 while the overall miscall rate was assessed 6e-04.

### Genetic structure and differentiation among localities

A pronounced pattern of population structure was observed among localities both from PCA and ADMIXTURE analyses. PCA showed evident clustering of individuals from the same locality (Supplementary Figure 1a). Although partly overlapping among the three habitat types (Supplementary Figure 1b), anadromous localities grouped together and distinct from river localities. Bayesian clustering analyses with the ADMIXTURE software for K = 48, 49, 50 showed that most localities could be assigned to a unique genetic cluster (Figure 2). The most admixed individuals mainly belong to two river populations (VIC and PEY) in the southern part of the study range in accordance with their geographic proximity. Moreover, ADMIXTURE analysis conducted on the seven anadromous localities displayed less pronounced structure, since K = 4, 5, 6 presented the lowest Cross-Validation errors. Overall, pairwise population differentiation (*F*_ST_) ranged from 0.01 to 0.33 (median = 0.17) (Supplementary Table 4) whereas mean pairwise *F*_ST_ per population ranged from 0.10 (VIC) to 0.24 (YAJ) (median = 0.17) (Table 2). Among lake populations, pairwise *F*_ST_ ranged from 0.05 to 0.33 (median = 0.19) while mean pairwise *F*_ST_ per population ranged from 0.13 (CON) to 0.25 (EKK) (median = 0.18). ^Among river populations, pairwise population *F*^_ST_ ranged from 0.02 to 0.17 (median = 0.10) while mean pairwise *F*_ST_ per population ranged from 0.08 (VIC) to 0.16 (FOR) (median = 0.09). Finally, anadromous populations displayed less pronounced pairwise *F*_ST_, with pairwise population *F*_ST_ ranging from 0.01 to 0.08 (median = 0.06), and mean pairwise *F*_ST_ per population ranging from 0.05 (AFE) to 0.08 (BWA) (median = 0.06) (Table 2). Together, these analyses confirmed a general pattern of genetic structure reflecting a situation of higher constrained gene flow and more pronounced genetic drift in freshwater (lake and riverine) than anadromous populations. Finally, there was no evidence of isolation by distance (IBD) among all populations (adj.R^2^ = 0.00, *P* = 0.536) and an extremely low (albeit significant) signal of IBD was observed when considering lake populations only (adj.R^2^ = 0.02, *P* = 7.69e-08), thus reflecting a highly constrained contemporary gene flow between resident populations.

**Figure 2.**
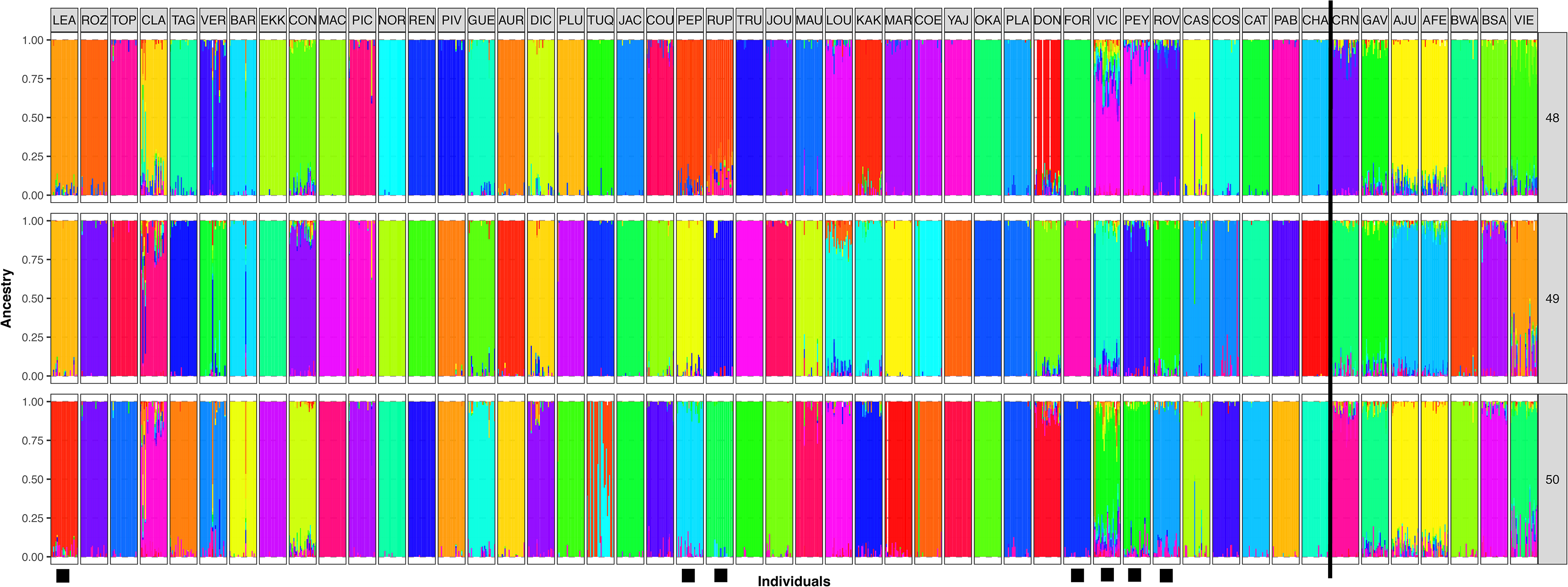
Admixture results for K = 48, 49 and 50. Localities are ordered according to geography (mainly from West to East). The solid black line delimitates the seven anadromous localities (right side of the line) and river localities are denoted by a black square above the locality.

**Table 2:** Population names and corresponding habitat type, differentiation estimates (Mean Fst across all pairs of localities, across lakes, across rivers and across anadromous localities), diversity (Π) within locality, proportion of polymorphic SNP, ratio of deleterious mutations and the ratio of deleterious mutations over ratio of polymorphic SNPs.

| NAME | Habitat | Latitude | Longitude | MeanFst | MeanFst<br>Lakes | MeanFst<br>Anadromous | MeanFst<br>River | $\Pi$ | Polymorphic<br>prop | ratio of<br>deleterious<br>mutat<br>ions | ratio of<br>deleterio<br>us<br>mutatio<br>ns over<br>ratio of<br>polymor<br>phic<br>SNPs |
| --- | --- | --- | --- | --- | --- | --- | --- | --- | --- | --- | --- |
| ROZ | lake | 54.827778 | -72.924722 | 0.165 | 0.179 | - | - | 0.00084033 | 0.40 | 0.89 | 2.21 |
| TOP | lake | 49.31327 | -79.19431 | 0.174 | 0.188 | - | - | 0.00081117 | 0.37 | 0.55 | 1.50 |
| CLA | lake | 48.30139 | -77.17167 | 0.118 | 0.131 | - | - | 0.00111753 | 0.52 | 0.69 | 1.31 |
| TAG | lake | 46.7094444 | -78.9969444 | 0.155 | 0.170 | - | - | 0.00099228 | 0.48 | 0.74 | 1.54 |
| VER | lake | 46.387222 | -78.311944 | 0.154 | 0.168 | - | - | 0.00092639 | 0.46 | 0.85 | 1.84 |
| BAR | lake | 46.6111005 | -76.6346647 | 0.223 | 0.234 | - | - | 0.00065355 | 0.38 | 0.75 | 1.94 |
| EKK | lake | 47.08944 | -75.94028 | 0.240 | 0.251 | - | - | 0.00060849 | 0.27 | 0.58 | 2.13 |
| CON | lake | 47.122834 | -75.009625 | 0.113 | 0.127 | - | - | 0.00105604 | 0.49 | 0.70 | 1.43 |
| MAC | lake | 46.9827 | -74.35416 | 0.199 | 0.209 | - | - | 0.00074846 | 0.33 | 0.79 | 2.37 |
| PIC | lake | 46.729494 | -74.870646 | 0.180 | 0.191 | - | - | 0.00083441 | 0.40 | 0.37 | 0.92 |
| NOR | lake | 46.105 | -75.346388 | 0.206 | 0.217 | - | - | 0.00071489 | 0.33 | 0.77 | 2.32 |
| REN | lake | 46.056704 | -74.486227 | 0.228 | 0.238 | - | - | 0.0007033 | 0.32 | 0.28 | 0.90 |
| PIV | lake | 46.49083 | -74.04472 | 0.210 | 0.219 | - | - | 0.00069805 | 0.34 | 0.79 | 2.34 |
| GUE | lake | 46.47638 | -73.86333 | 0.179 | 0.190 | - | - | 0.00080616 | 0.42 | 0.74 | 1.78 |
| AUR | lake | 46.464169 | -73.605553 | 0.150 | 0.165 | - | - | 0.00105706 | 0.47 | 0.68 | 1.45 |
| DIC | lake | 46.731985 | -73.215392 | 0.174 | 0.187 | - | - | 0.00080492 | 0.42 | 0.72 | 1.71 |
| PLU | lake | 47.126433 | -73.110941 | 0.172 | 0.183 | - | - | 0.00084626 | 0.37 | 0.36 | 0.95 |
| TUQ | lake | 47.457432 | -72.950555 | 0.208 | 0.220 | - | - | 0.00070549 | 0.36 | 0.70 | 1.94 |
| JAC | lake | 48.086446 | -74.598497 | 0.208 | 0.219 | - | - | 0.00071195 | 0.33 | 0.70 | 2.16 |
| COU | lake | 48.2663889 | -72.70444444 | 0.163 | 0.175 | - | - | 0.0008211 | 0.40 | 0.75 | 1.85 |
| TRU | lake | 49.94444 | -74.255 | 0.208 | 0.220 | - | - | 0.00069636 | 0.32 | 0.30 | 0.92 |
| JOU | lake | 49.810833 | -72.186111 | 0.164 | 0.175 | - | - | 0.00082429 | 0.40 | 0.60 | 1.53 |
| MAU | lake | 50.438057 | -71.713987 | 0.180 | 0.190 | - | - | 0.00075994 | 0.38 | 0.62 | 1.64 |
| LOU | lake | 49.135725 | -70.939472 | 0.173 | 0.181 | - | - | 0.00078416 | 0.42 | 0.68 | 1.63 |
| KAK | lake | 49.2188889 | -69.98 | 0.173 | 0.181 | - | - | 0.00078598 | 0.40 | 0.84 | 2.07 |
| MAR | lake | 48.651114 | -70.768069 | 0.162 | 0.172 | - | - | 0.00083358 | 0.38 | 0.74 | 1.94 |
| COE | lake | 48.67913 | -69.96453 | 0.168 | 0.178 | - | - | 0.00077003 | 0.39 | 0.73 | 1.85 |
| YAJ | lake | 50.218925 | -69.80186389 | 0.244 | 0.251 | - | - | 0.00057065 | 0.26 | 0.18 | 0.67 |
| OKA | lake | 50.371389 | -68.914444 | 0.211 | 0.218 | - | - | 0.00068645 | 0.31 | 0.75 | 2.38 |
| PLA | lake | 51.871485 | -68.166193 | 0.151 | 0.166 | - | - | 0.00093602 | 0.41 | 0.32 | 0.79 |
| DON | lake | 49.3191667 | -68.43083333 | 0.115 | 0.130 | - | - | 0.00102842 | 0.50 | 0.78 | 1.58 |
| CAS | lake | 48.06527778 | -68.30611111 | 0.178 | 0.197 | - | - | 0.00085543 | 0.42 | 0.74 | 1.75 |
| COS | lake | 48.15583333 | -68.70888889 | 0.160 | 0.176 | - | - | 0.00090888 | 0.41 | 0.71 | 1.71 |
| CAT | lake | 48.49694444 | -67.1525 | 0.167 | 0.186 | - | - | 0.00085484 | 0.39 | 0.70 | 1.77 |
| PAB | lake | 48.296711 | -64.924211 | 0.204 | 0.219 | - | - | 0.00075312 | 0.36 | 0.64 | 1.80 |
| CHA | lake | 50.617945 | -61.875805 | 0.179 | 0.197 | - | - | 0.0008645 | 0.40 | 0.60 | 1.52 |
| VIC | river | 45.47495 | -71.07359 | 0.104 | - | - | 0.078 | 0.00105413 | 0.51 | 0.89 | 1.74 |
| PEY | river | 46.2454 | -71.2384 | 0.120 | - | - | 0.090 | 0.00101316 | 0.47 | 0.79 | 1.69 |
| ROV | river | 46.53140296 | -70.57785488 | 0.113 | - | - | 0.087 | 0.00103096 | 0.47 | 0.75 | 1.59 |
| RUP | river | 51.033571 | -73.759052 | 0.145 | - | - | 0.095 | 0.00096375 | 0.40 | 0.45 | 1.15 |
| PEP | river | 51.39552778 | -72.97980333 | 0.147 | - | - | 0.100 | 0.00088995 | 0.42 | 0.86 | 2.02 |
| FOR | river | 45.37983 | -74.20815 | 0.197 | - | - | 0.159 | 0.00083612 | 0.37 | 0.75 | 2.03 |
| LEA | river | 57.918572 | -72.977104 | 0.152 | - | - | 0.108 | 0.00091061 | 0.48 | 0.77 | 1.63 |
| CRN | anadromous | 48.103889 | -66.282222 | 0.127 | - | 0.058 | - | 0.00097505 | 0.47 | 0.74 | 1.57 |
| GAV | anadromous | 49.083333 | -64.55 | 0.111 | - | 0.056 | - | 0.00104102 | 0.49 | 0.63 | 1.29 |
| AJU | anadromous | 49.476111 | -63.593611 | 0.123 | - | 0.054 | - | 0.00099781 | 0.48 | 0.67 | 1.40 |
| AFE | anadromous | 49.154167 | -62.715278 | 0.123 | - | 0.054 | - | 0.00099348 | 0.48 | 0.79 | 1.64 |
| BWA | anadromous | 50.216667 | -60.866667 | 0.155 | - | 0.076 | - | 0.00090556 | 0.44 | 0.67 | 1.51 |
| BSA | anadromous | 51.2 | -58.583333 | 0.132 | - | 0.055 | - | 0.00098168 | 0.48 | 0.71 | 1.47 |
| VIE | anadromous | 51.42194 | -57.94935 | 0.132 | - | 0.061 | - | 0.00094276 | 0.48 | 0.71 | 1.47 |

### Population genetic diversity

Overall, the median proportion of polymorphic SNPs per population was 40 %, ranging from 26 % (pop YAJ) to 52 % (pop CLA). The median value of observed heterozygosity was 0.14 and ranged from 0.12 (pop YAJ) to 0.15 (pop CLA). Over all 50 populations, the median value of average Π (considering both variable and non-variable sites) was 8.43E-04, ranging from 5.71E-04 (pop YAJ) to 1.11E-03 (pop CLA, Table 2 and Figure 3a). Median values of Π were 8.01E-04, 9.64E-04 and 9.82E-04 among lacustrine, riverine and anadromous populations, respectively. The mean Π was significantly lower among lake populations (mean Π _lakes_ = 8.16E-04) than among rivers (mean Π _rivers_ = 9.57E-04, *P_t_*_.test_ < 0.01) and among anadromous populations (mean Π _anadromous_ = 9.77E-04, *P_t_*_.test_ < 0.001). Mean Π among river populations was not significantly different from mean Π among anadromous populations (*P_t_*_.test_ = 0.31). No significant correlation was found between nucleotide diversity and latitude (adj.R^2^ = 0.00, *P* = 0.64, Figure 3b). However, a negative significant correlation was found between nucleotide diversity and altitude (*P* < 0.01, adj.R^2^ = 0.16, Figure 3c). A similar pattern was observed for *F*_ST_ which was not correlated to latitude (*P* = 0.34, adj.R^2^ = 0.00, Figure 3d), but was found to be positively correlated with altitude (*P* < 0.01, adj.R^2^ = 0.13, Figure 3e).

**Figure 3.**
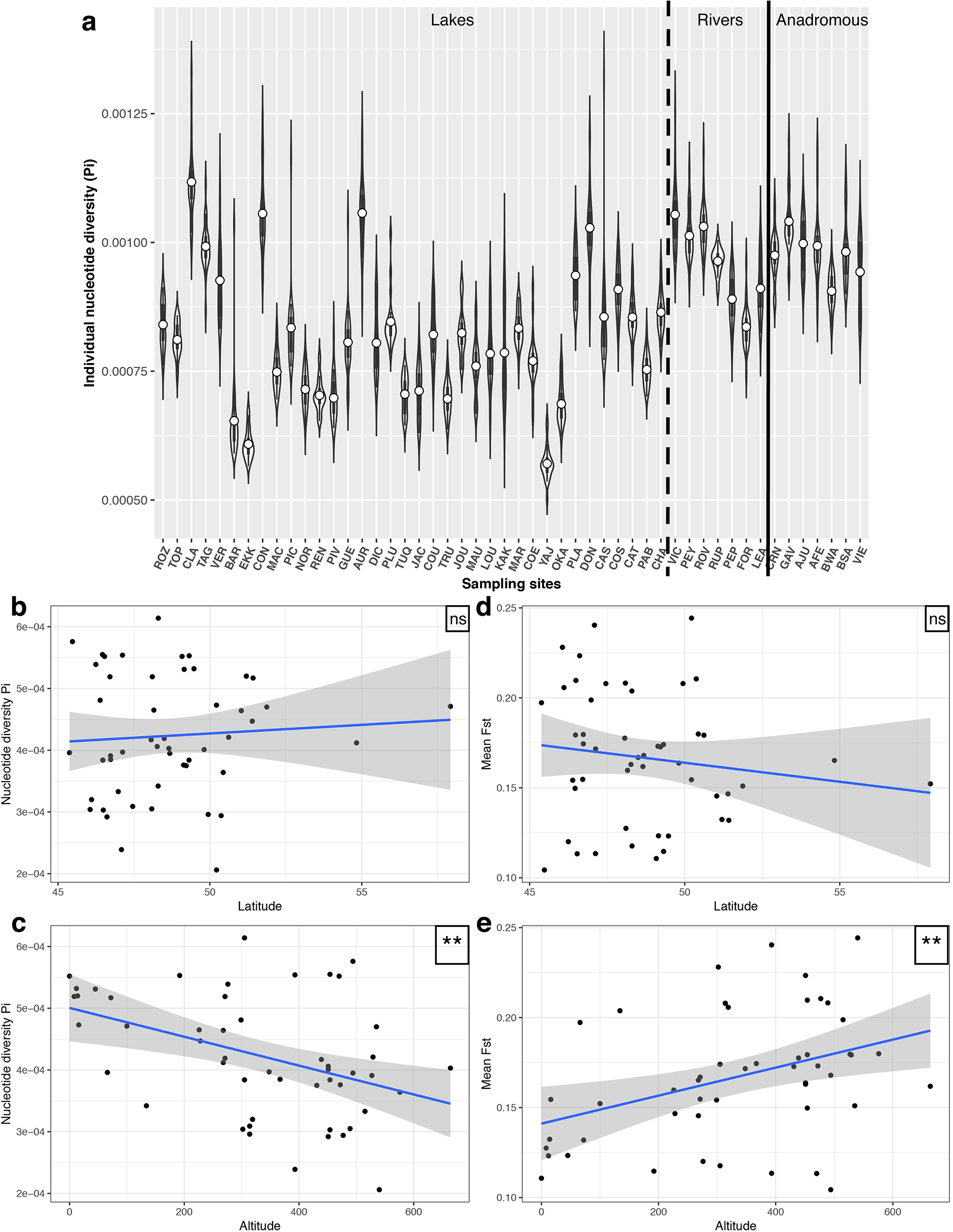
Relationships among genomic indices and environmental parameters. **(a)** Average individual nucleotide diversity across localities. Dashed line separates lakes from river localities, black line separates river from anadromous populations. Relationship between average nucleotide diversity and **(b)** latitude and **(c)** altitude, between Mean Fst and **(d)** latitude, and **(e)** altitude. “n.s.” denotes a non-significant relationship whereas “**” indicates a statistically significant relationship (p< 0.01).

### Putatively maladaptive (deleterious) mutations

Among the 4,729 genotyped loci (7,950 |SNPs), 1,475 SNPs had significant BLAST results against the Rainbow Trout transcriptome and were retained to assess synonymy. Synonymous substitutions were identified for 982 SNPs and nonsynonymous for 483 SNPs. Among the 483 nonsynonymous mutations, 289 were predicted to be neutral and 194 (40%) were putatively deleterious. The proportion of deleterious mutations within a given population varied from 0.15 (pop MAC and CHA) to 0.31 (pop GAV) (Table 2). The median of ratio of the proportion of deleterious mutations over the proportion of polymorphic SNPs was 1.64 and ranged from 0.67 (pop YAJ) to 2.383 (OKA, Table 2). This ratio was not significantly different between rivers and anadromous populations (t-tests: t_rivers∼ anadromous_ = 1.752, *P*_rivers∼ anadromous_ = 0.12) neither between lakes and rivers (t_lakes ∼ rivers_ = 0.15088, *P*_lakes ∼ rivers_ = 0.88), but it was significantly lower in anadromous populations (mean _anadromous_ = 1.48) compared to lakes (mean _lakes_ = 1.67, t_lakes ∼anadromous_ = 2.1692, *P*_lakes ∼ anadromous_ = 0.03,). Moreover, the average MAF of putative deleterious SNPs (mean _MAF deleterious_ = 0.14) was significantly lower than anonymous (mean _MAF anonymous_ = 0.16, t = −7.3895, *P* < 0.001), as well as non-synonymous but non-deleterious (mean _MAF nsnd_ = 0.20, t = −21.273, *P* < 0.001) and synonymous SNPs (mean _MAF synonymous_ = 0.22, t = −32.329, *P* < 0.001, Figure 4). Within putative deleterious SNPs no significant difference was observed between average MAF of habitat types (t_lakes ∼ rivers_ = 0.00, *P*_lakes ∼ rivers_ = 0.89, t_lakes ∼ anadromous_ = 1.514, *P*_lakes ∼ anadromous_ = 0.13, t_rivers∼ anadromous_ = 1.2653, *P*_rivers∼ anadromous_ = 0.21).

**Figure 4.**
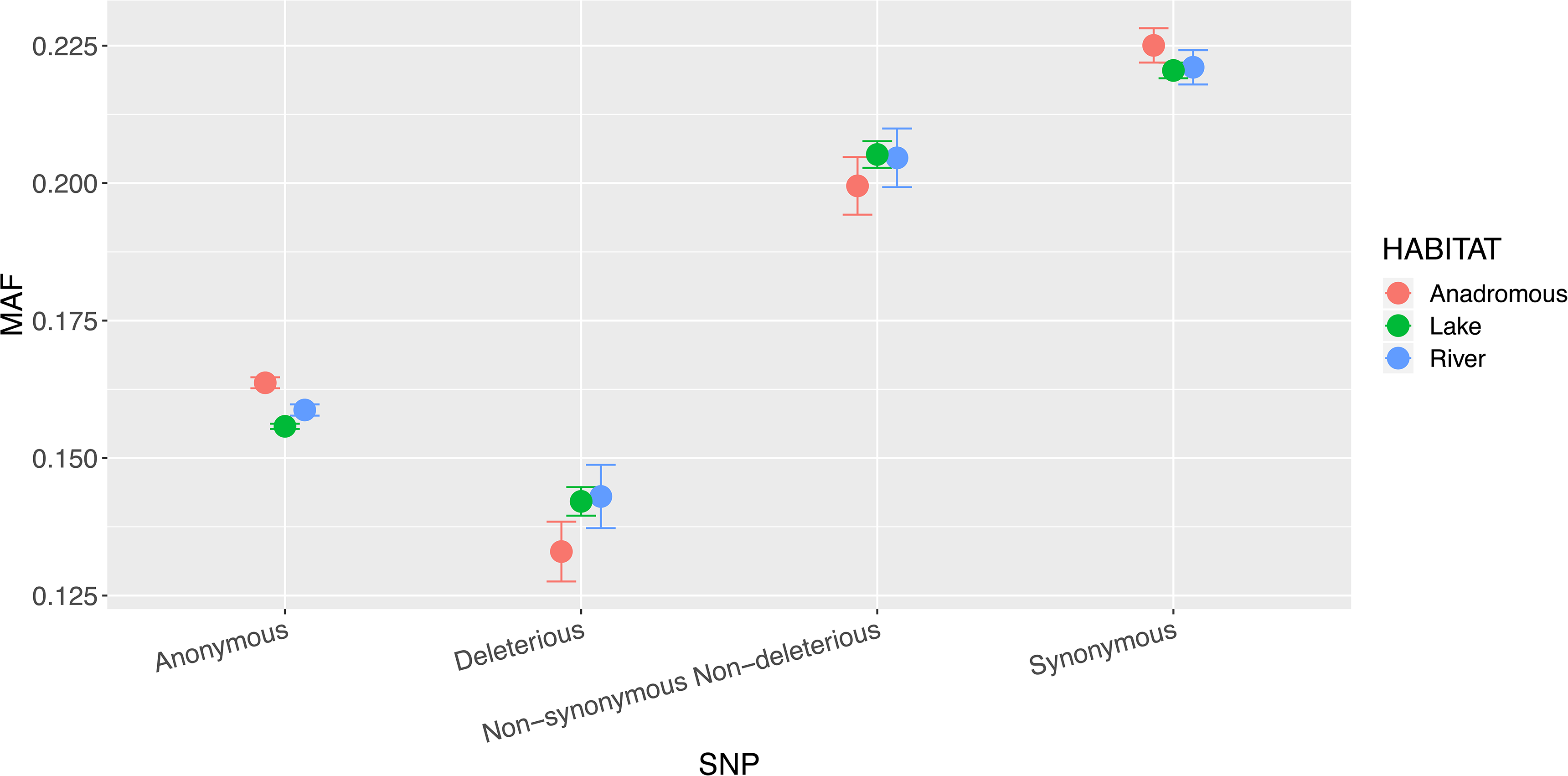
Average Minor Allele Frequency (MAF) across categories of SNPs (anonymous, deleterious, non-synonymous but non-deleterious and synonymous) within each category of habitat (anadromous in red, lakes in green and rivers in blue). Deleterious SNPs are significantly present in lower frequency than other SNPs but the difference of MAF between habitat types is not significant.

### Putatively adaptive mutations

Two analyses were performed to identify putatively adaptive mutations; first a test of SNPs associated with anadromy and then a test of association of SNPs with environmental variables among lacustrine populations only.

#### Anadromy

To correct for spatial auto-correlation, 27 axes were obtained from the db-MEM analysis, and among them seven were significant and therefore as a spatial component *via* an explanatory matrix. A total of 159 SNPs were found to be associated with anadromy when correcting for spatial autocorrelation (Supplementary Figure 2a), compared to 173 that were discovered without correction (Supplementary Figure 2b; both analyses with *P*_outlier_ < 0.001). Fifty-six SNPs were shared between these two lists and used as SNPs significantly associated with anadromy for subsequent analyses. After removing eight SNPs in common with putative deleterious mutations, we retained 48 loci significantly associated with anadromy (Supplementary Figure 3).

#### GEA in lacustrine populations

Among lacustrine populations, global model retained two out of the three environmental variables to be significantly associated with genetic variability (LowTmin and GDD). LowTmin was mostly associated with the second axis of the RDA which explained 4.8 % of the variation in allelic frequencies among lake populations (*P* = 0.002), while GDD was associated to the first axis of the RDA explaining 6.5% of the variation (*P* = 0.001, Supplementary Figure 4a). A total of 133 candidate SNPs (distributed across 125 loci) related to axis 1 represented a multi-locus set of SNPs associated with GDD while 159 candidate SNPs (distributed among 144 loci) related to axis 2 represented a multi-locus set of SNPs associated with LowTmin (*P*_outlier_ < 0.001; Supplementary Figure 4b). After removing SNPs in common with putative deleterious mutations, we retained respectively 119 and 137 loci significantly associated with GDD and LowTmin loci (*P*_outlier_ < 0.001; Supplementary Figure 3) and identified as putative beneficial mutations linked to environmental conditions.

### Genomic positions and gene ontology

Among all 4729 loci genotyped across all 50 populations, 3585 were successfully mapped against the Arctic Charr genome. On average, among both putative adaptive and maladaptive variants, 82 % (ranging from 81 % for GDD to 84 % for markers linked to anadromy) of the defined candidate loci mapped to the genome while an average of 57 % for putatively adaptive and 71 % for putatively maladaptive mutations were located in annotated regions (Table 3 and Figure 5). Among the DNA sequences successfully annotated, 65 % (over all annotations) corresponded to transposons (Supplementary Table 5), thus reflecting a statistically significant enrichment for transposable elements among adaptive and maladaptive markers relative to the reference transcriptome (*P* < 0.001). More than half (54%) of those transposons corresponded to Tc1-like class transposons (Supplementary Table 5). The proportion of transposons was higher for DNA sequences associated with environmental variables (73 % LowTmin, 76 % for GDD and 83 % Anadromy) than for DNA sequences including putatively deleterious variants (60 %). Other than transposons, annotated DNA sequences included genes representing multiple biological functions potentially involved in local adaptation. For SNPs associated with anadromy, two genes (Type 1 inositol 1,4,5-trisphosphate receptor and Na(+)/Ca(2+)-exchange protein 1) involved in homeostasis were found as well as one related to cardiac muscle differentiation and one associated with neuron differentiation (Supplementary Table 5). Biological functions such as immune response, homeostasis, male gonad development and oogenesis characterized genes associated to temperature (LowTmin). One gene involved in immune response (DEAD box protein 58) was also discovered in genomic regions associated with GDD as well as other functions including adult locomotion behavior and muscle organ development. Three genes involved in fatty acid metabolic process (Fat/vessel-derived secretory protein, Lupus nephritis-associated peptide 1 and Palmitoyl-CoA oxidase) were also discovered within genes associated with GDD. Two genes also involved in fatty acid metabolism were found as putatively deleterious (ELOVL fatty acid elongase 1 and Cytochrome P450 2D14). Other biological functions discovered for putative deleterious genes comprised neurogenesis, cardiac muscle cell differentiation, apoptosis, circadian regulation, embryonic development, lifespan, ossification and immune response (Supplementary Table 5).

**Figure 5.**
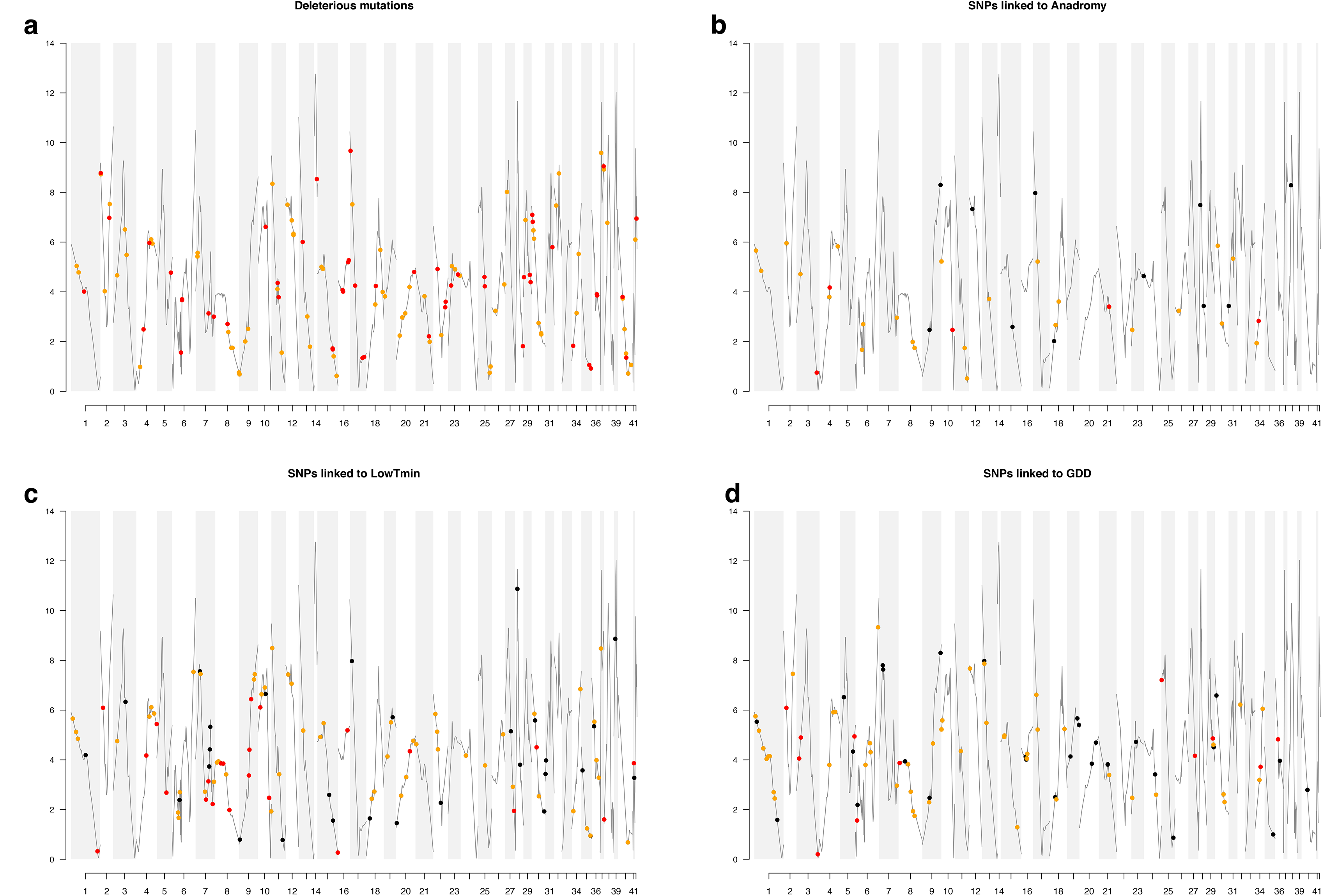
Inference of recombination rate along the Brook Charr linkage groups (in cM/ Mbp). Both putative deleterious SNPs and SNPs associated with environmental variables are represented on their genomic localization. Black dots denote SNPs into non-coding regions, orange dots represent SNPs falling into a Transposable Element (TE) and red dots correspond to SNP into coding regions with biological functions other than TEs.

**Table 3:** Number of deleterious and beneficial variants that have been detected, mapped to the genome and annotated to any biological function.

|  | **Total nb of**<br>**SNPs** | **Total nb of**<br>**loci** | **Nb of**<br>**mapped SNPs** | **Nb_of**<br>**mapped loci** | **Percent of**<br>**mapped loci** | **Nb of**<br>**annotated**<br>**SNPs** | **Nb of**<br>**annotated**<br>**loci** | **Percent of**<br>**annotated**<br>**loci** |
| --- | --- | --- | --- | --- | --- | --- | --- | --- |
| **GDD** | 133 | 125 | 109 | 101 | 80.80% | 72 | 67 | 53.60% |
| **LowTmin** | 159 | 144 | 134 | 117 | 81.25% | 95 | 85 | 59.03% |
| **Anadromy** | 56 | 51 | 49 | 43 | 84.31% | 36 | 30 | 58.82% |
| **Deleterious** | 177 | 177 | 148 | 148 | 83.62% | 126 | 126 | 71.19% |

### Effect of recombination rate on the genomewide distribution of putative adaptive and maladaptive mutations

Variation of inferred recombination rate along the Brook Charr reconstructed genome (based on the 3582 mapped positions) ranged from 1.98 cM/ Mbp to 12.68 cM/ Mbp (Figure 5). The average recombination rate per chromosome ranged from 1.98 cM/ Mbp (CH 41) to 10.22 cM/ Mbp (CH 14), with minimal recombination rate ranging from 0.07 cM/ Mbp (CH 1) to 7.95 cM/Mbp (CH 14), and maximal recombination rate ranging from 3.86 cM/ Mbp (CH 41) to 12.68 cM/ Mbp (CH 14). The highest standard deviation of recombination rate within chromosome was in CH 17 (3.15 cM/ Mbp) while CH 16 (0.53 cM/ Mbp) harbored the lowest standard deviation. Regions comprising putative deleterious mutations did not show a different mean recombination rate than for the rest of SNPs (mean R _del_ = 4.31 ± 2.25, mean R _non-del_ = 4.46 ± 2.37, *P* = 0.48, Figure 6). The same pattern was observed for SNPs associated with anadromy (mean R _ana_ = 3.94 ±2.03, mean R _non-ana_ = 4.46 ± 2.37, *P* = 0.13, Figure 6) and SNPs associated with environmental variables among lacustrine populations (mean R _GDD_ = 4.37 ± 1.8, mean R _non-GDD_ = 4.46 ± 2.37, *P* = 0.66; mean R _LowTmin_ = 4.22 ± 2.12, mean R _non-LowTmin_ = 4.46 ± 2.37, *P* = 0.33, Figure 6). Finally, for non-synonymous variants, no correlation was found between Provean scores and recombination rate (adj.R^2^ = 0.00, *P* = 0.68, Figure 7).

**Figure 6.**
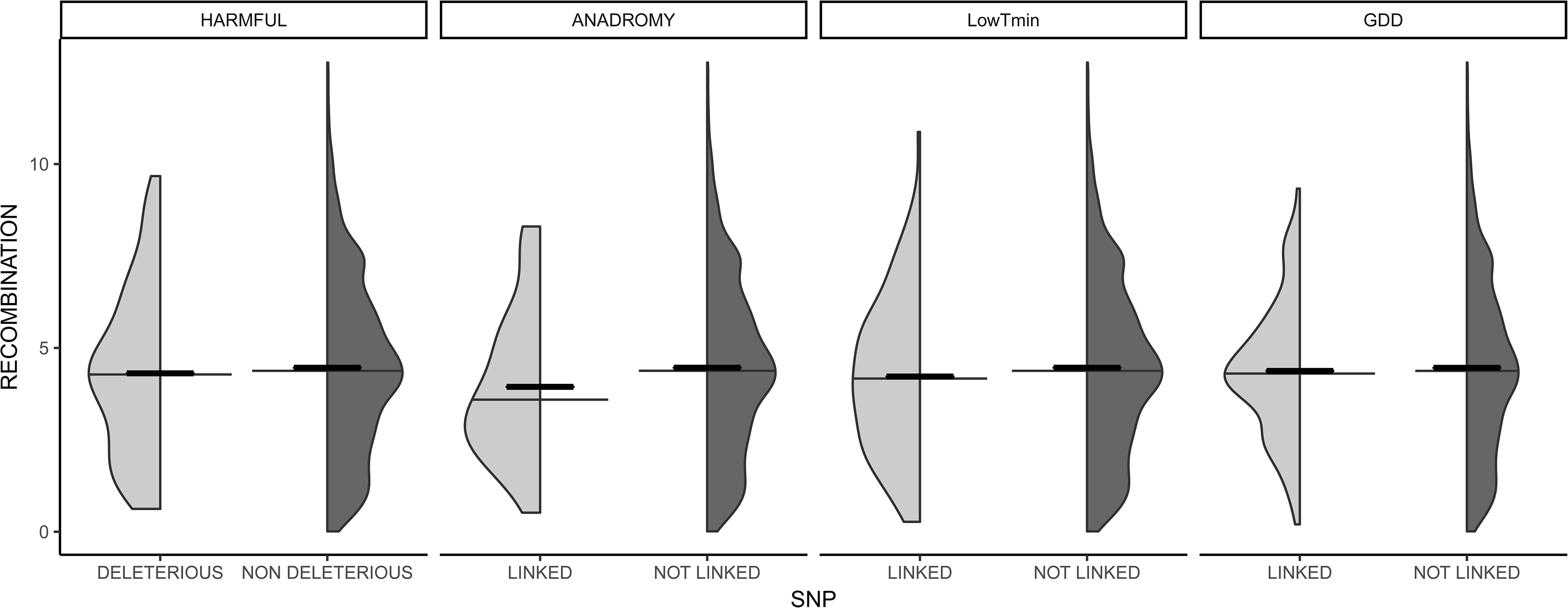
Boxplot of recombination rate (in cM/Mbp) across SNPs defined as putatively deleterious *versus* those that are non-deleterious and SNPs associated to environmental variable *versus* those not linked to the respective environmental variable

**Figure 7.**
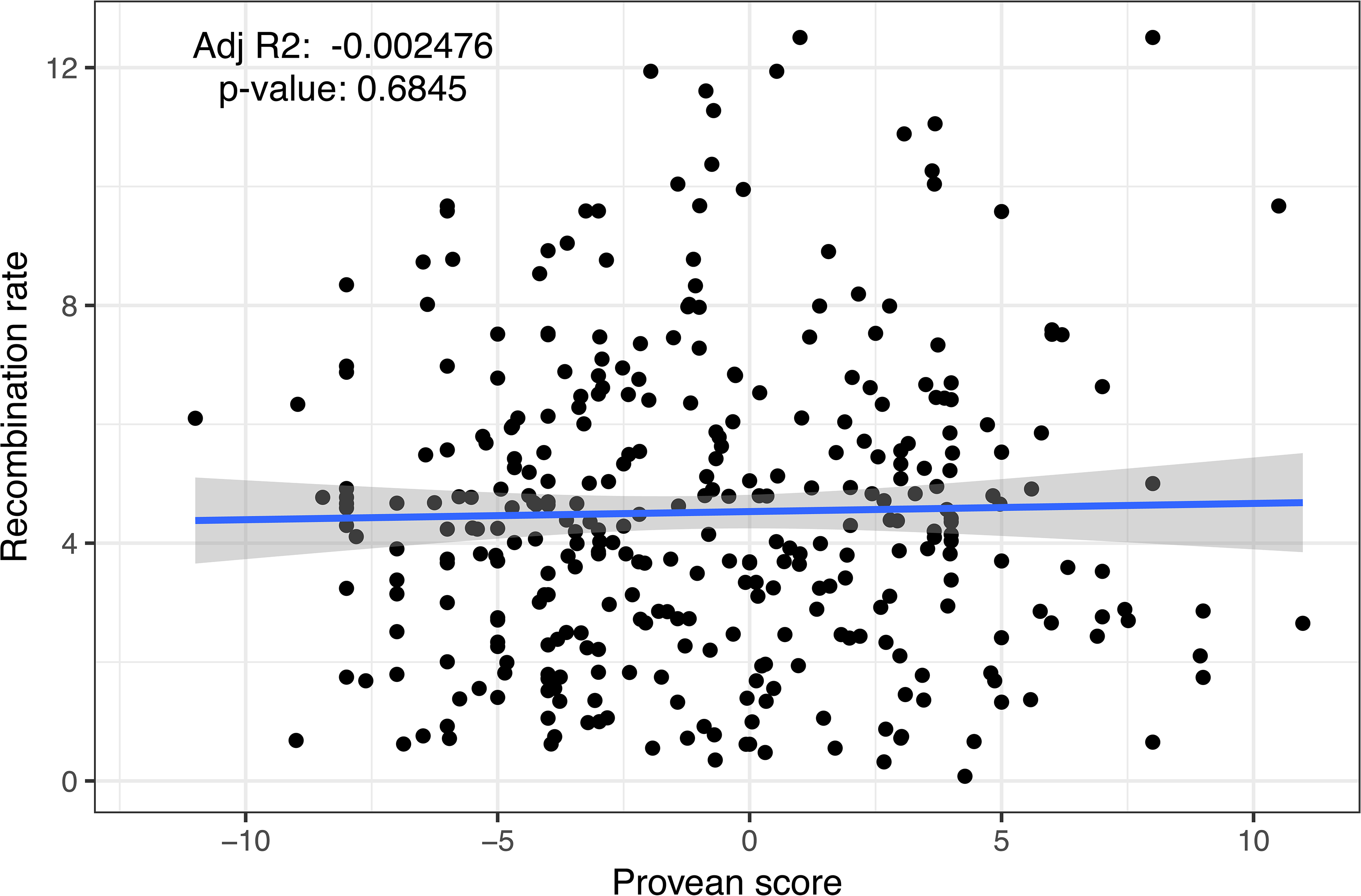
Relationship between recombination rate (in cM/Mbp) and deleteriousness (i.e. Provean score equal or below than −2.5 the variants is considered to be putatively harmful and above −2.5 is considered to be neutral)

## Discussion

The goal of this study was to investigate the relative importance of demography and selection in the accumulation of both putative adaptive and maladaptive mutations in small populations. Firstly, we detected a pronounced genetic structure among Brook Charr lacustrine populations along with a reduced genetic diversity compared with river and anadromous populations. The absence of IBD combined with the observed relationship between genetic diversity, differentiation and altitude suggest that colonization history has played an important role in the accumulation of both neutral and (mal)adaptive variants in this system. As expected in such systems with an overriding role of demographic processes on genetic variation, a non-negligible accumulation of deleterious mutations was found in Brook Charr. Moreover, a negligible role of linked selection was revealed by the absence of relationship found between deleteriousness and recombination rate. Finally, functional genes found to be associated with environmental variables and/or the anadromy supports a significant effect of selection in maintaining local adaptation in the face of the apparent prevailing role of genetic drift in shaping genetic variation in Brook Charr populations.

### Pronounced genetic structure and reduced genetic diversity in lacustrine populations

Brook Charr is considered among the most highly structured animal species (Ward et al 1994), and our results confirm it. The pronounced level of structuration observed in our Brook Charr samples have been reported (i) at large scale in which genetic variance is partitioned among major drainages or regions associated with distinct refugial origins (Perkins et al 1993; Danzmann et al. 1998), and (ii) at a finer geographical scale (few kilometers) (Angers and Bernatchez 1998; Angers et al 1999; Hébert et al 2000). More pronounced structuration among lake than river and anadromous populations highlights the fact that gene flow among Brook Charr populations depends on habitat availability favoring dispersion. The absence or very limited evidence of isolation by distance we observed also reflects the highly constrained contemporary gene flow (albeit to a lesser extent in anadromous populations) and as well as a departure from migration-drift equilibrium. Such a scenario was previously proposed to explain genetic differentiation among Brook Charr populations in Maine, USA in which IBD exists but decreases in the most recently colonized northern populations (Castric et al 2001). Our results corroborate and strengthen this pattern.

The reduced genetic diversity in lakes compared with rivers and anadromous populations further confirmed the pattern of a highly restricted contemporary gene flow. Moreover, the reduction of genetic diversity with altitude along the increase of differentiation strengthens the impact of colonization history on the observed genetic variability (Castric et al 2001). Indeed, high-altitude populations are expected to be more physically isolated (and therefore more genetically differentiated), either because of increased probability of physical barriers to gene flow (*e.g.* impassable waterfalls) and/or due to more pronounced founder effects, assuming that the number of colonists decrease with altitude (Sagarin and Gaines 2002).

### Accumulation of maladaptive (deleterious) mutations

According to population genetics theory, the accumulation of deleterious mutations depends on the relative roles of multiple factors including mutation rate, demographic history and selection. In accordance with theoretical expectations, we observed that putative deleterious mutations were on average observed at smaller frequencies than the rest of the variants within a given population. Overall, neither the accumulation of putative deleterious variants nor their predicted level of deleteriousness by Provean were correlated with recombination rate, suggesting that (i) the role of putative linked selection may be negligible, and that (ii) genetic drift might be the major factor explaining the randomly potential harmful mutations genomic distribution. Globally, albeit to a lower extent than previously reported in isolated lacustrine populations of Lake Trout (60 % of nonsynonymous were putatively deleterious, Perrier, Ferchaud et al. 2017) we observed a significant accumulation of deleterious mutations in Brook Charr populations (39 % of nonsynonymous were putatively deleterious). For comparison, the percentage of nonsynonymous sites estimated to be deleterious in previous studies ranges from 3% in bacterial populations (Hughes 2005) up to 80% in the human population (Fay et al 2001). Such relative abundant levels of deleterious mutations in Brook and Lake Trout might be attributed to their colonization history and to their small effective population sizes, which are known to cause the accumulation of deleterious variants overtime (Benazzo et al 2017; Grossen et al 2019). The lower occurrence of deleterious mutations in Brook Charr compared to Lake Trout could be imputable to differences in their life history. Lake Trout is a top predator and a large, long-lived fish restricted to lacustrine populations generally harboring very small effective population sizes (Wilson and Mandrack 2003), whereas Brook Charr is a much smaller fish on average with a shorter generation time harboring both resident and anadromous populations (Scott and Crossman 1973). Lake Trout is almost entirely restricted to lacustrine environment whereas Brook Charr is also commonly found is streams and rivers, therefore providing to this species more opportunity for connectivity among populations compared to Lake Trout. In particular, anadromous populations which are less genetically differentiated are therefore likely less affected by random genetic drift effect due to their dispersal abilities, also revealed a tendency to harbor a lower accumulation of putative deleterious mutations than lake or river populations. Admittedly however, our assessment of deleteriousness must be interpreted cautiously. Indeed, we used the functionality of a given mutation based on the degree of conservation of an amino acid residue across species (Choi et al 2012). However, as in previous studies that applied this approach, those assumptions have not been experimentally validated, and there is no causal relationship available between the occurrence of these putative deleterious mutations and reduced fitness. Despite these caveats, those methods to assess the accumulation of deleterious have recently been applied to estimate genetic load in human populations (Peischl et al 2017) and domesticated species (Renaut and Rieseberg 2015; Zhou et al 2016) and have been performed to validate theoretical predictions by empirical observations (Balick et al 2015). Moreover, a very recent infatuation for such approaches has emerged for managing wild populations (Benazzo et al 2017; Perrier, Ferchaud et al 2017; Ferchaud, Laporte et al 2018; Zhu et al 2018; Grossen et al 2019). Indeed, in addition to support the process of adaptation, the management decisions must adequately account for maladaptation as a potential outcome and even as a tool to bolster adaptive capacity to changing conditions (Derry et al 2019).

### What are the putative adaptive and maladaptive mutations?

Among annotated regions identified for the putative adaptive and maladaptive mutations, a large proportion (65%) shows a statistically significant enrichment of transposable elements. Transposable Elements (TEs) are short DNA sequences with the capacity to move between different genomic locations in the genome (Bourque et al 2018). This ability may generate genomic mutations in many different ways, from subtle regulatory mutations to large genomics rearrangements (Casacuberta and Gonzalez 2013). The body of evidence regarding the role of such genomic variants in evolutionary biology is growing (Dion-Côté and Barbash 2017) and recently TEs evolution have been suggested to contribute to adaptation to climate change (Rey et al. 2016, Lerat et al 2019, Shrader et al 2019). First, TEs generate mutations actively by inserting themselves in new genomics locations and passively by acting as targets of ectopic recombination (recombination between homologous sequences that are not at the same position on homologous chromosomes). Second, some TEs contain regulatory sequences that can affect the structure and the expression of the host genes (Chuong et al 2016; Elbarbary et al 2016). Salmonid genomes are known to contain a large quantity of transcribed mobile elements (Krasnov et al 2005). In particular, TEs are suspected to play a major role in reshaping genomes and playing an important role in genome evolution (DeBoer et al 2007, Rodriguez and Arkhipova 2018), especially following genome duplications (Ewing 2015). Salmonids have undergone a whole genome duplication event around 60 Mya (Crête-Lafrenière, Weir and Bernatchez 2012) followed by rediploidization events (*i.e.* return to stable diploid states) through genome rearrangements such as fusions and fissions (Sutherland et al 2016). The majority of TEs we identified in Brook Charr belongs to Tc1-like class transposons (54 %). Interestingly, the expression profiles of Tc1-like transposons (the most widespread mobile genetic elements) have been shown to be strongly correlated with genes implicated in defense response, signal transduction and regulation of transcription in salmonid fish (Krasnov et al 2005). On the other hand, we also identified TEs associated with putative maladaptive variants (55% of annotated putative deleterious variants), as expected considering that previous studies showed that TEs can have profoundly deleterious consequences on organisms (Goodier 2016). Previous studies have also evidenced the effect of TEs on deleteriousness in other salmonids. In common garden environments, hybrid breakdown in Lake whitefish (*Coregonus clupeaformis*) has been associated with TEs reactivation caused by a genomic shock (genomic incompatibility) and associated hypomethylation in hybrids, leading to a malformed phenotype and aneuploidy (Dion-Côté et al. 2014; Dion-Côté et al. 2017; Laporte et al. in revision). Overall, this highlights the importance to increase our understanding of such structural variants in small populations.

Other biological functions identified for the putative beneficial and harmful mutations are relevant in the context of maladaptation (for deleterious ones) or adaptation to anadromy and temperature. Two putative targets of selection identified linked with anadromy are related to homeostasis and one is related with cardiac muscle differentiation. This observation strengthens the fact that migration might be a strong agent of selection as already suggested by other studies in salmonids (Pritchard et al 2018) and particularly in the genus *Salvelinus* (Moore et al 2017). Fish and fisheries researchers have yet to widely acknowledge the possible role Growing Degree-Day (GDD) having another potentially powerful selective agents acting on fish growth and development. Neuheimer and Taggart (2007) showed that fish length-at-day was a strong linear function of the GDD metric that explains > 92% of growth. However, our study is the first (to our knowledge) showing an association between GDD and genetic variation possibly underlying growth. Indeed, outliers putatively associated with GDD are located in genomic regions with biological functions related to fatty acid metabolic process, organ development and locomotion behavior which all relevant in growth process. Lastly, genomic regions putatively associated with temperature also corresponded to potentially relevant biological functions such as immune system, as observed previously in Lake Trout (Perrier, Ferchaud et al 2017). Interestingly, adaptation to local temperature regime and bacterial community prevailing in the natal river has been supported through large-scale variation in MHC diversity, suggesting the influence of pathogen-driven balancing selection on MHC diversity in Atlantic salmon (Dionne et al 2007) and underline the importance of MHC standing genetic variation for facing pathogens in a changing environment (Dionne et al 2009).

## Conclusion

The relative roles of demography and selection are a key in understanding the process of the maintenance of adaptive and maladaptive variation in small populations. Particularly, understanding the factors that cause harmful mutations to increase in frequency in a genome will facilitate prediction of genetic load in current populations, and in this way contribute to improve conservation practices, for instance by providing a rationale for genetic rescue. Indeed, increasing empirical evidence suggests that recently fragmented populations with reduced population sizes may receive demographic benefits from gene flow beyond the addition of immigrant individuals through genetic rescue (Frankham 2015). However, most of empirical studies conducted on wild populations to date are incomplete because of the difficulty of rigourously assess the impact of detected genotype-environment or genotype-phenotype associations on individual fitness (but see Wells et al 2019). As such, our study adds to the increasing realisation that in a context of a rapidly changing environment, the risk of maladaptation must be considered in planning conservation strategies.

## Acknowledgments

We thank biologists and technicians of the Ministère des Forêts, de la Faune et des Parcs du Québec (MFFP) for their implication in the project and their field assistance as well as all the outfitters who contributed to the sampling. We also thank Guillaume Côté and Clément Rougeux for discussion since the preliminary result of this study. We are grateful to the team of the genomic analyses platform of IBIS (Institut de Biologie Intégrative des Systèmes) at Laval University. This research was funded by the MFFP, the Canadian Research Chair in Genomics and Conservation of Aquatic Resources, Ressources Aquatiques Québec (RAQ) as well as by a Strategic Project Grant from the Natural Sciences and Engineering Research Council of Canada (NSERC) to L. Bernatchez, D. Garant and P. Sirois.

## Authors’ contribution

Conceived the study: L.B., I.T. and A-L.F. Logistics to collect samples: A-L.F., I.T. and C.H. Laboratory work: B.B. and D.B. Bioinformatics analyses: A-L.F. and E.N. Statistical analyses and writing the manuscript: A-L.F., M.Le. and M.La. All co-authors critically revised and contributed to edit the manuscript and approved the final version to be published.

## Supplementary material

**Supplementary Fig.1** Principal Component Analysis (PCA) produced on 7,950 SNPs of 1,193 Brook Charr where individuals **(a)** were colored by locality and **(b)** by habitat type.

**Supplementary Fig.2** Redundancy analyses produced on 7,950 SNPs of 50 Brook Charr populations explained by habitat with **(a)** correction by geography and **(b)** without correction.

**Supplementary Fig.3** Venn diagram of the number of SNPs defined as putatively deleterious, associated with anadromy, LowTmin and/or Growth Day Degree.

**Supplementary Fig.4** Redundancy analyses on 7,950 SNPs of 36 Brook Charr lakes. Scaling highligths (a) population variation, and b) outlier SNPs.

**Supplementary Table 1** Population names and corresponding environmental variables used in this study. Localities are ordered to their habitat type and subsequently according to geography (mainly from West to East), same order is used all along the figures and tables of the manuscript.

**Supplementary Table 2** Detailed methods, options and values for each filter used to identify SNPs for this study on unstocked Brook Charr populations in Québec, Canada.

**Supplementary Table 3** Details of the number of SNPs or Genotypes retained for each filter applied on the genomic dataset.

**Supplementary Table 4** Average pairwise Fst estimates among the 50 Brook Charr populations

**Supplementary Table 5** Annotation results, either sum up for all categories or detailed for each category (deleterious, associated with anadromy, LowTmin and GDD)

