## Supplementary figures and images for "Adaptive and maladaptive genetic diversity in small populations; insights from the Brook Charr (*Salvelinus fontinalis)* case study"

### Supplementary Fig.1

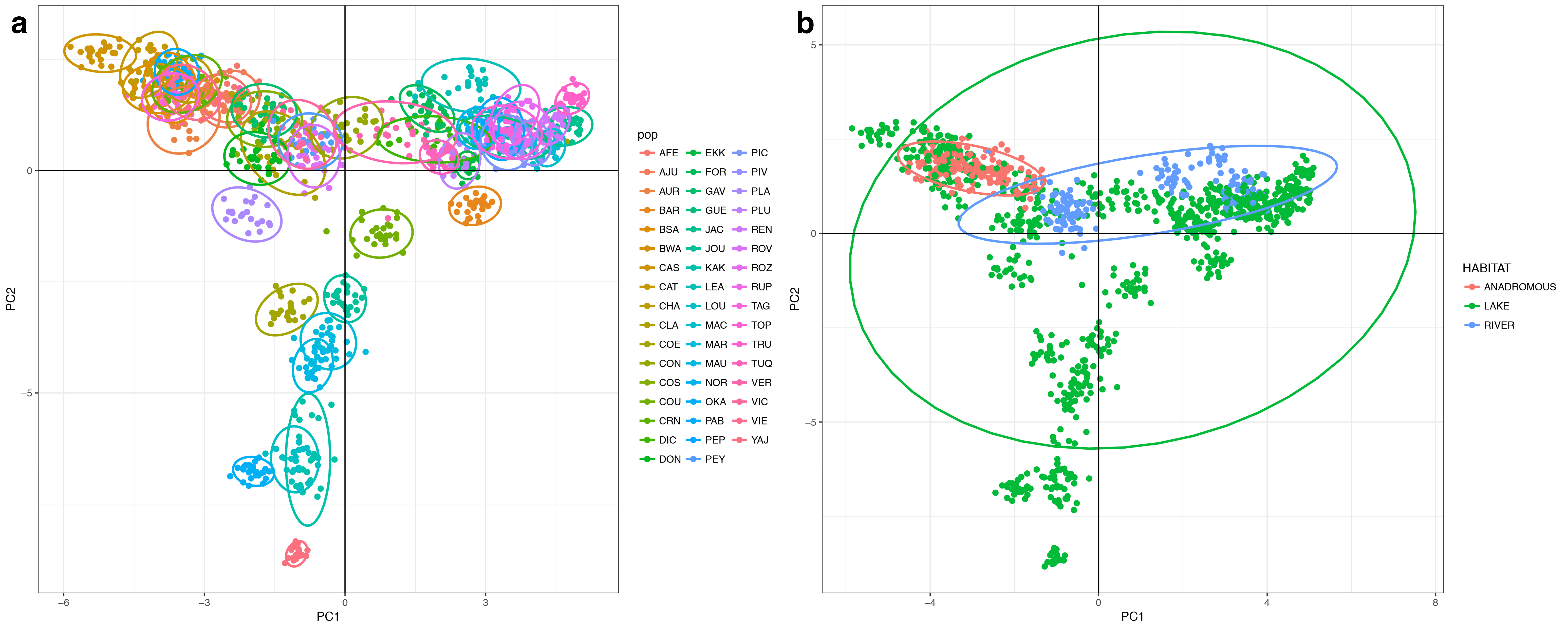

### Supplementary Fig.2

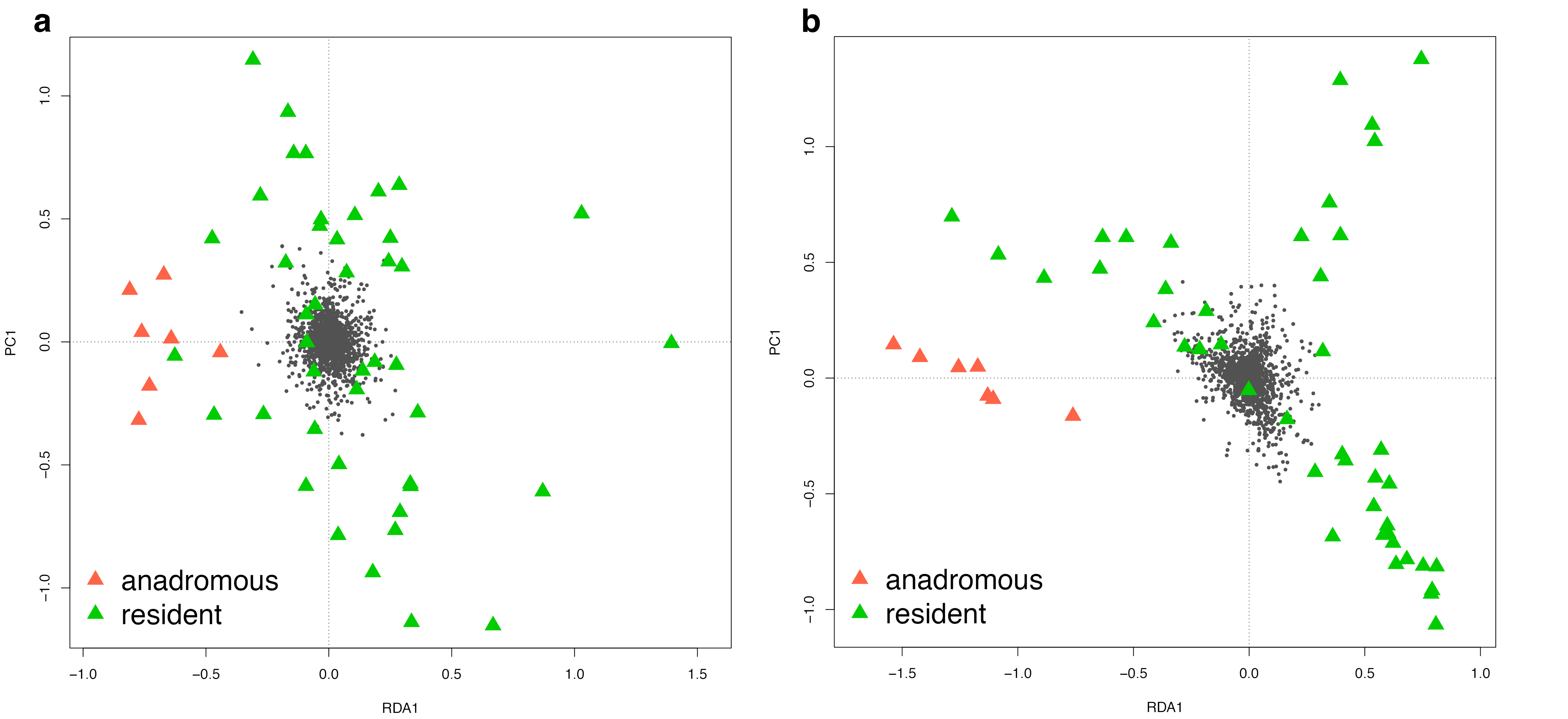

### Supplementary Fig.3

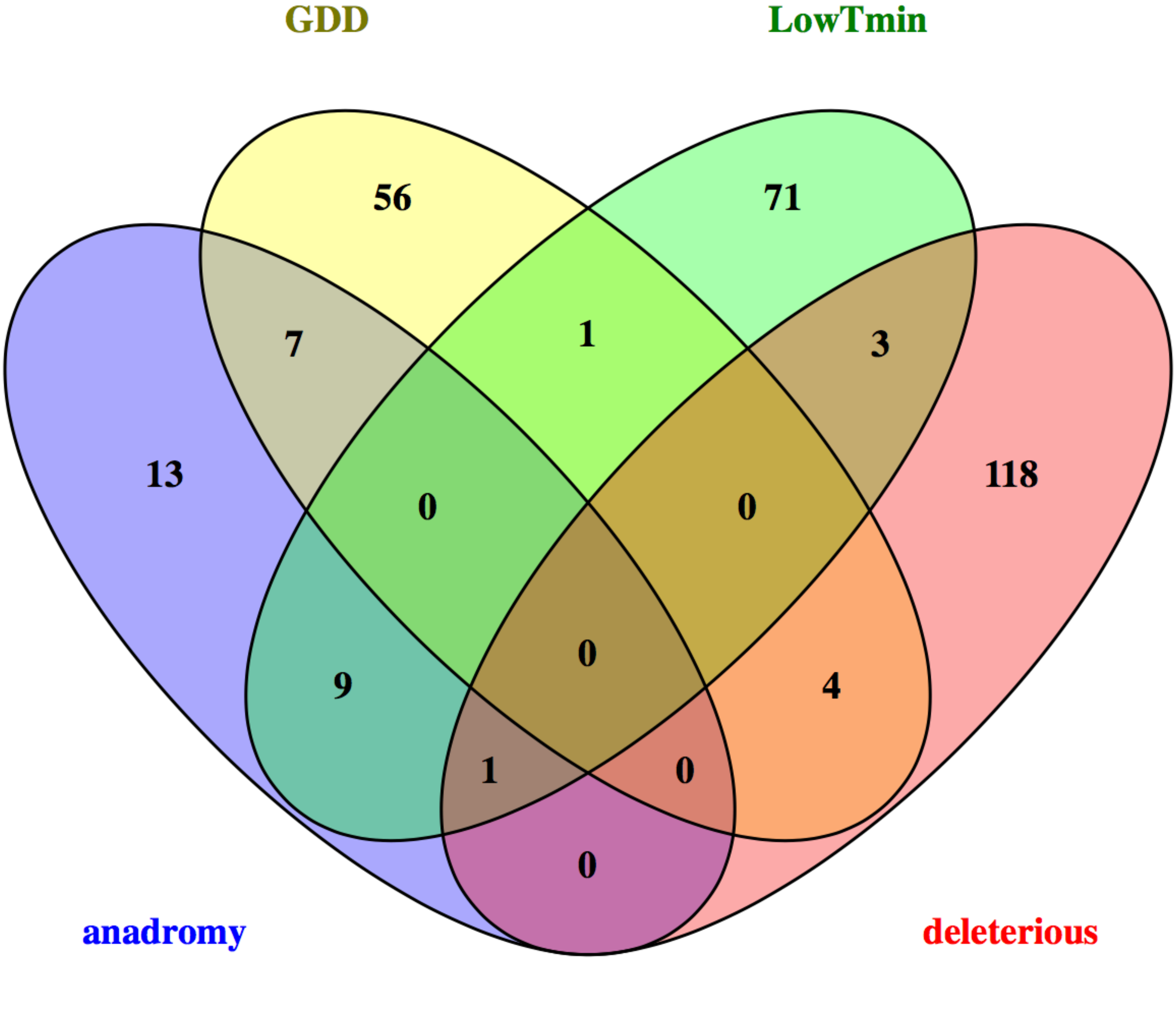

### Supplementary Fig.4

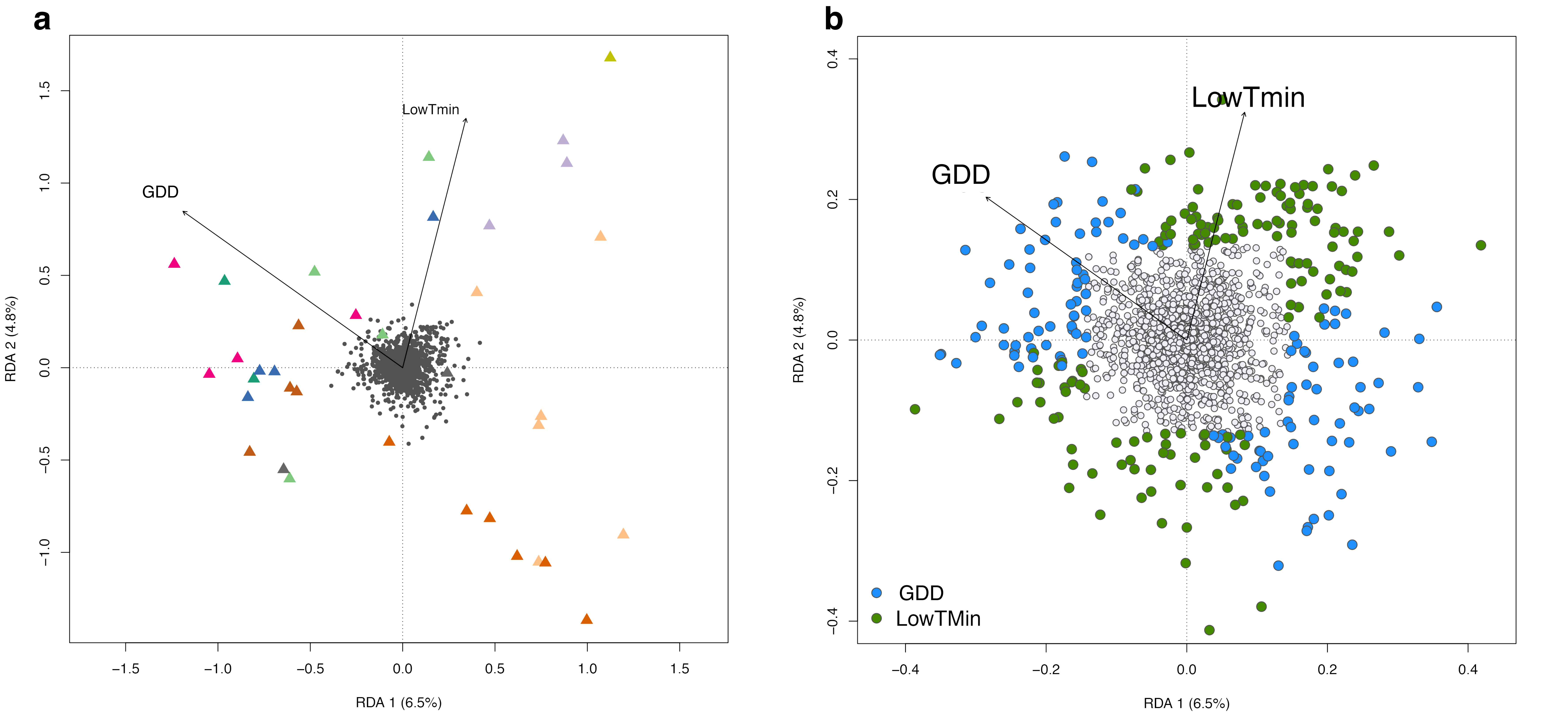
